# Specific hypersensitive response-associated recognition of new apoplastic effectors from *Cladosporium fulvum* in wild tomato

**DOI:** 10.1101/127746

**Authors:** Carl H. Mesarich, Bilal Ökmen, Hanna Rovenich, Scott A. Griffiths, Changchun Wang, Mansoor Karimi Jashni, Aleksandar Mihajlovski, Jérôme Collemare, Lukas Hunziker, Cecilia H. Deng, Ate van der Burgt, Henriek G. Beenen, Matthew D. Templeton, Rosie E. Bradshaw, Pierre J.G.M. de Wit

**Affiliations:** Laboratory of Phytopathology, Wageningen University, Droevendaalsesteeg 1, 6708 PB Wageningen, the Netherlands; Laboratory of Molecular Plant Pathology, Institute of Agriculture & Environment, Massey University, Private Bag 11222, Palmerston North 4442, New Zealand; Bio-Protection Research Centre, New Zealand; College of Chemistry and Life Sciences, Zhejiang Normal University, Jinhua, Zhejiang 321004, People’s Republic of China; Department of Plant Pathology, Iranian Research Institute of Plant Protection, Agricultural Research, Education and Extension Organization, P.O. Box 19395-1454, Tehran, Iran; Institute of Fundamental Sciences, Massey University, Private Bag 11222, Palmerston North 4442, New Zealand; Breeding & Genomics/Bioprotection Portfolio, the New Zealand Institute for Plant & Food Research Limited, Mount Albert Research Centre, Auckland 1025, New Zealand; Centre for BioSystems Genomics, P.O. Box 98, 6700 AB Wageningen, the Netherlands.

**Keywords:** Effectoromics, *Cf* immune receptor genes, apoplastic effectors, antimicrobial proteins, *Cladosporium fulvum*, *Solanum lycopersicum* (tomato)

## Abstract

Tomato leaf mould disease is caused by the biotrophic fungus *Cladosporium fulvum*. During infection, *C. fulvum* produces extracellular small secreted protein (SSP) effectors that function to promote colonization of the leaf apoplast. Resistance to the disease is governed by *Cf* immune receptor genes that encode receptor-like proteins (RLPs). These RLPs recognize specific SSP effectors to initiate a hypersensitive response (HR) that renders the pathogen avirulent. *C. fulvum* strains capable of overcoming one or more of all cloned *Cf* genes have now emerged. To combat these strains, new *Cf* genes are required. An effectoromics approach was employed to identify wild tomato accessions carrying new *Cf* genes. Proteomics and transcriptome sequencing were first used to identify 70 apoplastic *in planta*-induced *C. fulvum* SSPs. Based on sequence homology, 61 of these SSPs were novel or lacked known functional domains. Seven, however, had predicted structural homology to antimicrobial proteins, suggesting a possible role in mediating antagonistic microbe−microbe interactions *in planta*. Wild tomato accessions were then screened for HR-associated recognition of 41 SSPs using the *Potato virus X*-based transient expression system. Nine SSPs were recognized by one or more accessions, suggesting that these plants carry new *Cf* genes available for incorporation into cultivated tomato.

## INTRODUCTION

Leaf mould disease of tomato (*Solanum lycopersicum*) is caused by the biotrophic Dothideomycete fungal pathogen *Cladosporium fulvum* (syn. *Passalora fulva* and *Fulvia fulva*) (Thomma et al., 2005). The fungus likely originated in South America, the centre of origin for tomato (Jenkins, 1948), with the first disease outbreak reported in South Carolina, USA, during the late 1800s (Cooke, 1883). *C. fulvum* now occurs worldwide, but is primarily a problem in greenhouse and high-tunnel environments, where tomato plants are exposed to both moderate temperatures and high relative humidity. Disease symptoms are typified by pale green to yellow spots on the adaxial leaf surface, as well as white to olive-green patches of mould on the abaxial leaf surface that turn brown upon sporulation. In the late stages of disease development, this sporulation is often associated with leaf wilting and partial defoliation, which, in severe infections, can cause death of the plant (Thomma et al., 2005).

During infection (i.e. in a compatible interaction), *C. fulvum* exclusively colonizes the tomato leaf apoplast, where it grows in close contact with surrounding mesophyll cells (Thomma et al., 2005). This colonization is promoted through a collection of virulence factors, termed effector proteins, which the fungus secretes into the apoplastic environment (e.g. Laugé et al., 1997). To date, 13 *C. fulvum* effectors have been identified, and the genes encoding these proteins have been cloned (Bolton et al., 2008; Joosten et al., 1994; Laugé et al., 2000; Luderer et al., 2002a; Mesarich et al., 2014; Ökmen et al., 2013; Stergiopoulos et al., 2012; van den Ackerveken et al., 1993; van Kan et al., 1991; Westerink et al., 2004). The majority (11 of 13) are small secreted proteins (SSPs) of less than 300 amino acid residues in length with: (i) an amino (N)-terminal signal peptide for secretion into the tomato leaf apoplast; and (ii) four or more cysteine (Cys) residues following their signal peptide cleavage site. An intrinsic virulence function has been determined for three of the 11 SSP effectors. The first of these, Avr2, which lacks a known functional domain, targets and inhibits at least four Cys proteases of tomato (Rcr3, Pip1, aleurain and TDI-65) to prevent the degradation of *C. fulvum* proteins (Krüger et al., 2002; Rooney et al., 2005; Shabab et al., 2008; van Esse et al., 2008). The second, Avr4, possesses a carbohydrate-binding module family domain (CBM_14; PF01607) that binds chitin present in the cell wall of *C. fulvum* to protect against hydrolysis by basic plant chitinases (van den Burg et al., 2004, 2006; van Esse et al., 2007). The third, Ecp6, possesses three lysin motif domains (LysM; PF01476) that function to perturb chitin-triggered immunity (Bolton et al., 2008; de Jonge et al., 2010; Sánchez-Vallet et al., 2013). More specifically, two of the LysM domains cooperate to sequester chitin fragments released from the cell wall of invading hyphae, and in doing so, outcompete host chitin immune receptors for the binding of chitin fragments (Sánchez-Vallet et al., 2013). The third LysM domain has been proposed to perturb chitin-triggered immunity through interference with the host chitin immune receptor complex (Sánchez-Vallet et al., 2013).

Despite their roles in virulence, the same effectors can also be an Achilles’ heel for *C. fulvum*. In particular accessions of tomato, these effectors or their modulated targets can be directly or indirectly recognized, respectively, as invasion patterns (IPs) by corresponding Cf immune receptors to trigger immune responses that render the pathogen avirulent (Cook et al., 2015; de Wit et al., 2009; Wulff et al., 2009b). In these incompatible interactions, the main output of the immune system is the hypersensitive response (HR), a localized form of cell death that arrests growth of the pathogen at the infection site (Heath, 2000). So far, 10 of the 11 *C. fulvum* SSP effectors, specifically Avr2, Avr4, Avr4E, Avr5, Avr9, Ecp1, Ecp2-1, Ecp4, Ecp5 and Ecp6, are known to be recognized as IPs in tomato accessions with the corresponding Cf immune receptors Cf-2.1/Cf-2.2, Cf-4, Cf-4E, Cf-5, Cf-9, Cf-Ecp1, Cf-Ecp2-1, Cf-Ecp4, Cf-Ecp5 and Cf-Ecp6, respectively (de Wit et al., 2009; Thomma et al., 2011). All *Cf* immune receptor genes cloned to date encode receptor-like protein (RLP) cell surface receptors that possess extracytoplasmic leucine-rich repeats (eLRRs), a transmembrane domain, and a short cytoplasmic tail (Dixon et al., 1996, 1998; Jones et al., 1994; Panter et al., 2002; Takken et al., 1999; Thomas et al., 1997). Several studies suggest that the eLRRs are responsible for the direct or indirect recognition of *C. fulvum* effector proteins in the tomato leaf apoplast (Seear and Dixon, 2003; van der Hoorn et al., 2001a; Wulff et al., 2001, 2009a).

It was determined early on that wild *Solanum* species and landraces are a rich source of resistance against *C. fulvum*. Indeed, all cloned *Cf* immune receptor genes are derived from wild *Solanum* species or landraces, with *Cf-2.1*/*Cf-2.2, Cf-9*/*Cf-9DC* and *Cf-9B* from *Solanum pimpinellifolium* (Dixon et al., 1996; Jones et al., 1994; Panter et al., 2002; van der Hoorn et al., 2001b), *Cf-4* and *Cf-4E* from *Solanum habrochaites* (Takken et al., 1999; Thomas et al., 1997), and *Cf-5* from the landrace *Solanum lycopersicum* var. *cerasiforme* (Dixon et al., 1998). Based on this knowledge, *Cf* immune receptor genes were introgressed from wild *Solanum* species and landraces into cultivated tomato by breeders over several decades (Kerr and Bailey, 1964 and references therein). While largely effective, intensive year-round cultivation of these plants has led to the emergence of natural *C. fulvum* strains capable of overcoming one or more of all cloned *Cf* immune receptor genes (Hubbeling, 1978; Iida et al., 2015; Laterrot, 1986; Li et al., 2015). Several types of sequence modification have been shown to occur in *IP* effector genes that permit the evasion of *Cf* immune receptor-mediated resistance by *C. fulvum*. These are: (i) gene deletion; (ii) the insertion of a transposon-like element (gene disruption); (iii) single nucleotide polymorphisms (SNPs) that result in non-synonymous amino acid substitutions; and (iv) nucleotide insertions or deletions (indels) that result in frame-shift mutations (Stergiopoulos et al., 2007). To combat strains capable of overcoming existing resistance specificities, new *Cf* immune receptor genes need to be identified for incorporation into cultivated tomato.

Laugé et al. (2000) hypothesized that “any stable, extracellular protein produced by a pathogen during colonization is a potential avirulence factor [IP]”. With this in mind, and given that all cloned *Cf* immune receptor genes encode an RLP, we set out to identify wild tomato accessions carrying new *Cf* immune receptor genes corresponding to apoplastic *in planta*-induced SSPs (ipiSSPs) of *C. fulvum* using effectoromics. Effectoromics is a powerful high-throughput functional genomics approach that uses effectors or effector candidates to probe plant germplasm collections for corresponding immune receptors (Domazakis et al., 2017; Du and Vleeshouwers, 2014; Vleeshouwers and Oliver, 2014). Notably, this approach, which is based on the HR-associated recognition of effectors or effector candidates, has already proven to be successful for the identification of wild accessions and breeding lines of *Solanum* carrying *Cf* immune receptor genes corresponding to known effectors of *C. fulvum*. In a pioneering study by Laugé et al. (1998), 21 *S. lycopersicum* lines originating from early *C. fulvum* resistance breeding programmes were screened for their ability to recognize Ecp2-1 using the *Potato virus X* (PVX)-based transient expression system (Hammond-Kosack et al., 1995; Takken et al., 2000), as well as by leaf injection with purified Ecp2-1 protein. Four lines, which have the same *S. pimpinellifolium* ancestor, recognized Ecp2-1, indicating for the first time that tomato carries an immune receptor gene corresponding to this effector (*Cf-Ecp2-1*) (Laugé et al., 1998).

In a follow-up study by Laugé et al. (2000), 28 *S. lycopersicum* breeding lines, many of which also have an *S. pimpinellifolium* ancestor, were screened for their ability to recognize purified Ecp1, Ecp2-1, Ecp3 (amino acid sequence not yet known), Ecp4 or Ecp5 protein. Four lines recognized Ecp2-1, while two different lines recognized Ecp3 and Ecp5, respectively (Laugé et al., 2000). In the same study, a collection of 40 different *S. pimpinellifolium* accessions were also screened for their ability to recognize the same five effectors, as well as Avr4 and Avr9, using the PVX-based transient expression system. Three different accessions recognized Ecp1, Ecp2-1 and Ecp3 (purified protein), respectively, while two recognized Ecp4, three recognized Ecp5, and six recognized Avr9 (Laugé et al., 2000). Again, this study indicated for the first time that tomato carries immune receptor genes corresponding to Ecp3 (*Cf-Ecp3*), Ecp4 (*Cf-Ecp4*) and Ecp5 (*Cf-Ecp5*) (Laugé et al., 2000). Three known *C. fulvum* effectors have since been shown to be recognized by wild tomato accessions through infiltration of purified protein, specifically Ecp6 in *S. lycopersicum* (Thomma et al., 2011), as well as Avr4 and Avr9 in *S. pimpinellifolium* (Kruijt et al., 2005; van der Hoorn et al., 2001b).

As a starting point for our effectoromics approach, we used proteomics and transcriptome sequencing to identify 70 apoplastic ipiSSPs of *C. fulvum*. This set of 70 is made up of all 11 known SSP effectors of this fungus, as well as 59 *C. fulvum* candidate effectors (CfCEs). We screened 41 of these ipiSSPs for HR-associated recognition by wild tomato accessions using the PVX-based transient expression system. A total of nine ipiSSPs, renamed as extracellular proteins (Ecps), were recognized by one or more of 14 wild tomato accessions, suggesting that these plants carry new *Cf* immune receptor genes available for incorporation into cultivated tomato.

## RESULTS

### Proteomics and transcriptome sequencing identify 70 apoplastic ipiSSPs of *C. fulvum*

Liquid-chromatography–tandem mass spectrometry (LC–MS/MS) was used to identify fungal peptides corresponding to SSPs present in intercellular washing fluid (IWF) samples of compatible *C. fulvum*–tomato (*S. lycopersicum* cv. Heinz [H]-Cf-0) interactions. Here, SSPs are defined as those proteins of less than 300 amino acid residues in length with a predicted N-terminal signal peptide, but without a predicted glycophosphatidylinositol (GPI) anchor modification site, one or more transmembrane domains, a carboxyl (C)-terminal endoplasmic reticulum (ER) retention (H/KDEL)/retention-like (XXEL) signal, or sequence homology to enzymes. Using this approach, 297 unique fungal peptides were mapped to 75 SSPs of *C. fulvum* (Table S1 and Information S1). Based on pre-existing RNA-Seq transcriptome sequencing data from a compatible *C. fulvum* strain 0WU–*S. lycopersicum* cv. H-Cf-0 interaction at 4, 8 and 12 d post-inoculation (dpi), as well as from *C. fulvum* strain 0WU grown *in vitro* in potato-dextrose broth (PDB) or Gamborg B5 liquid media at 4 dpi (Mesarich et al., 2014), 70 of the 75 apoplastic SSPs (∼93.3%) were deemed to be encoded by *in planta*-induced genes (Tables 1 and S1).

**Table 1.**
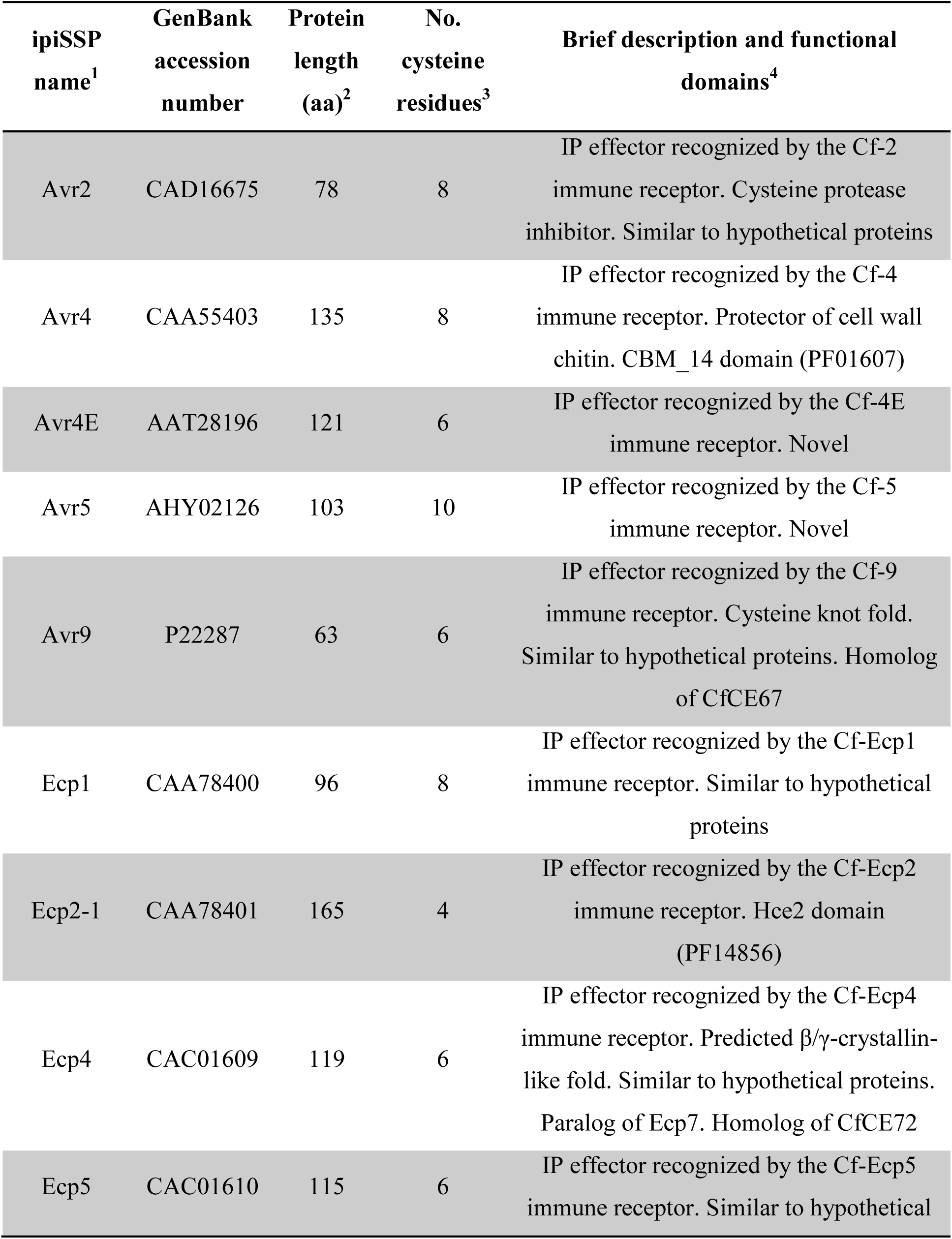
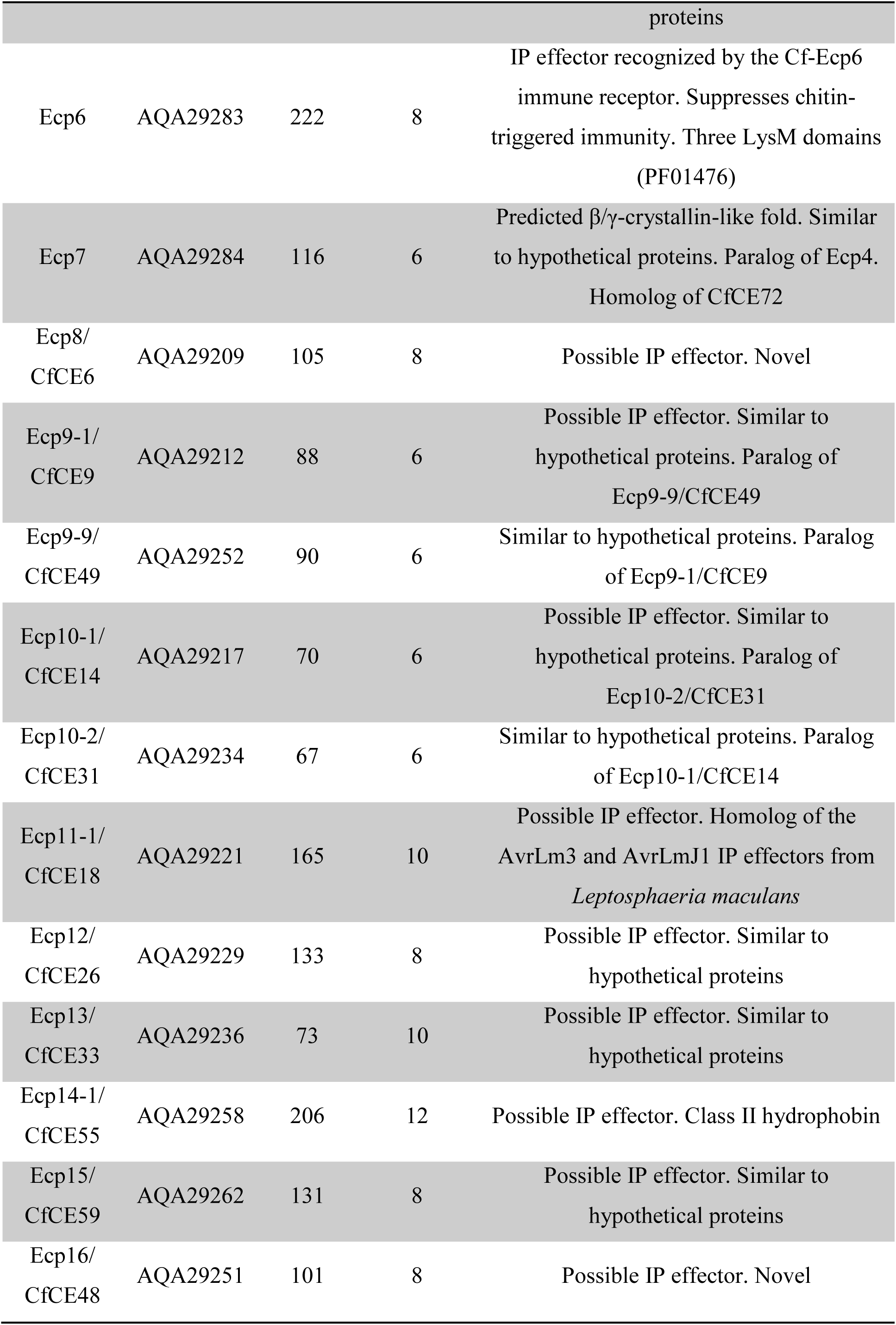
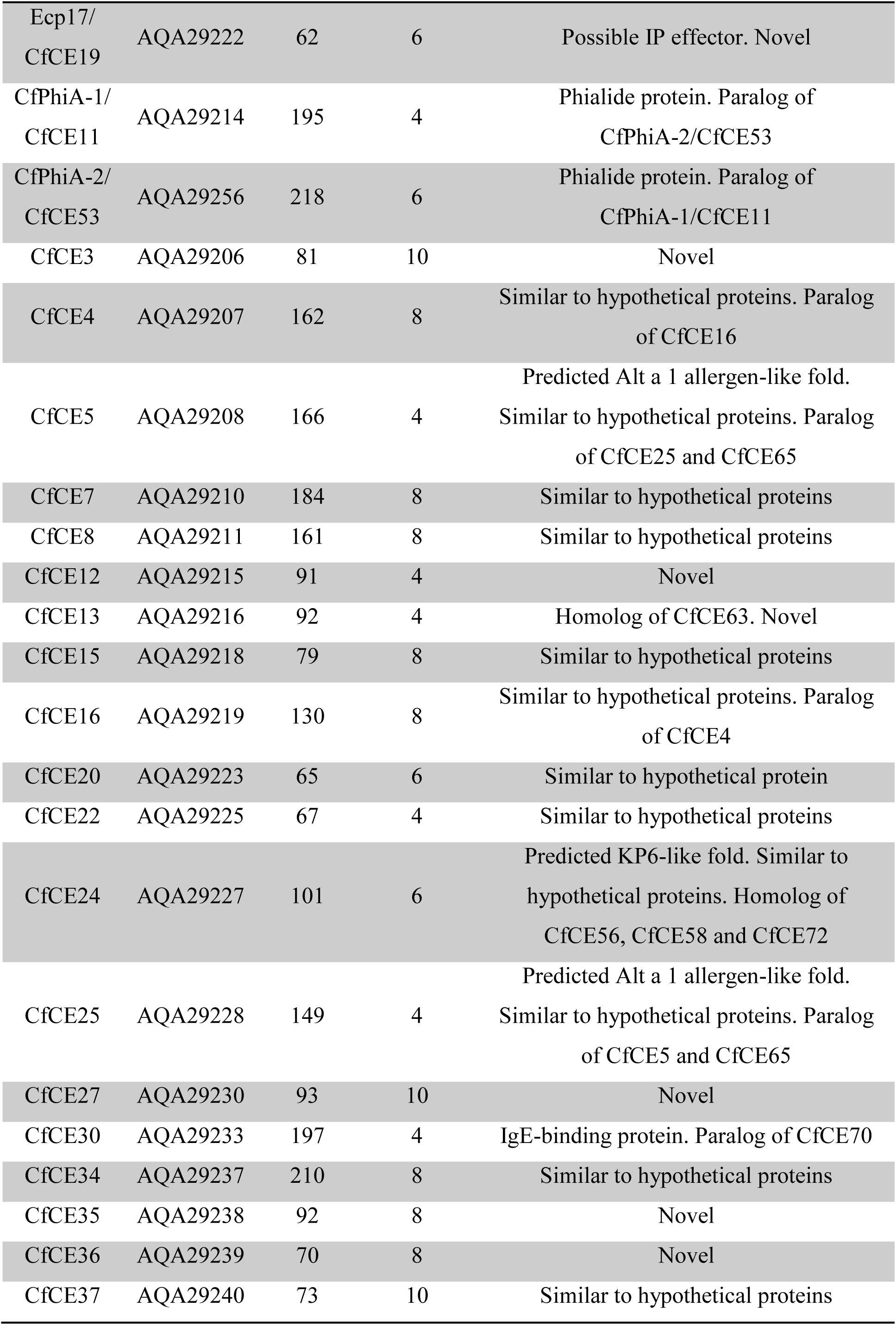
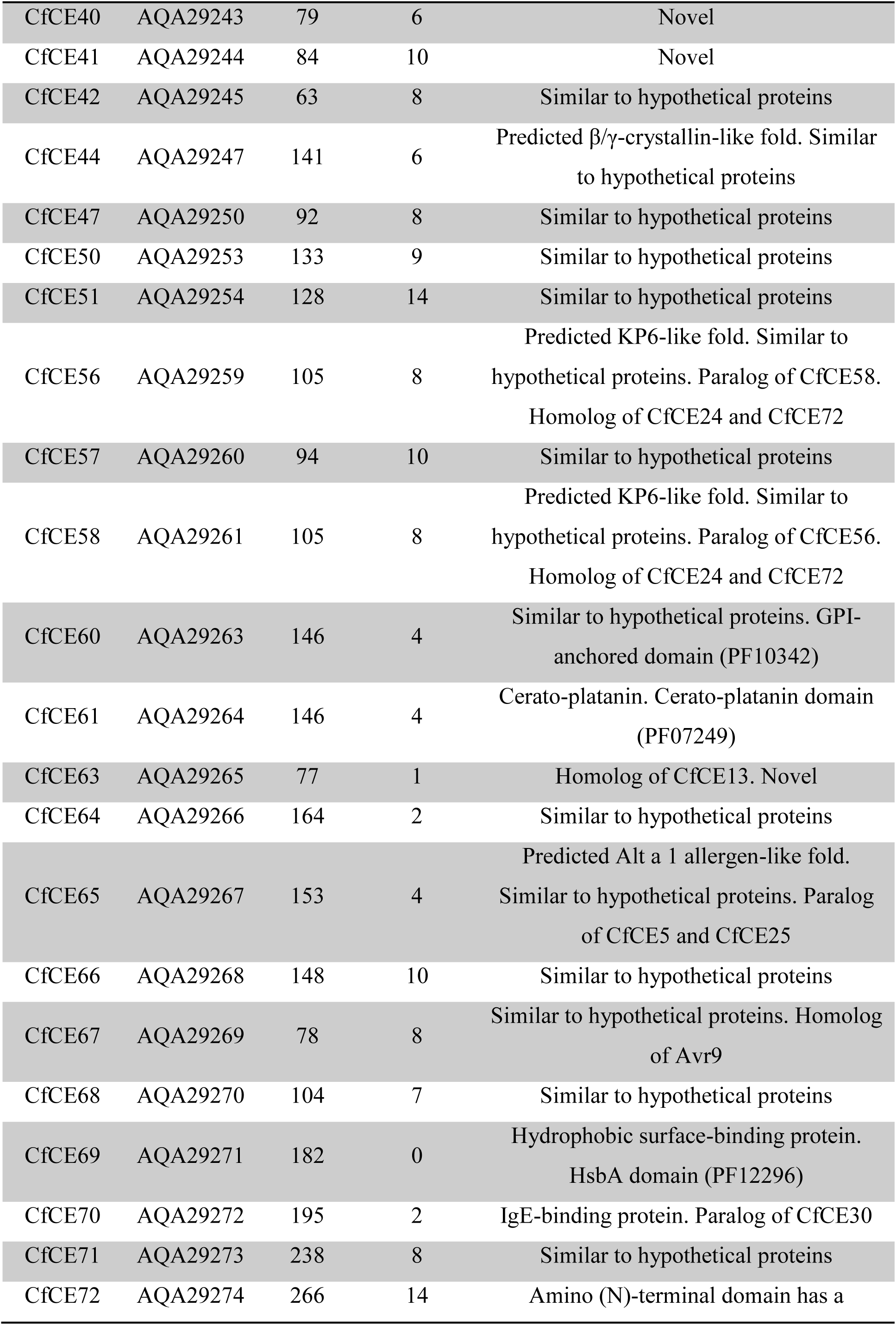
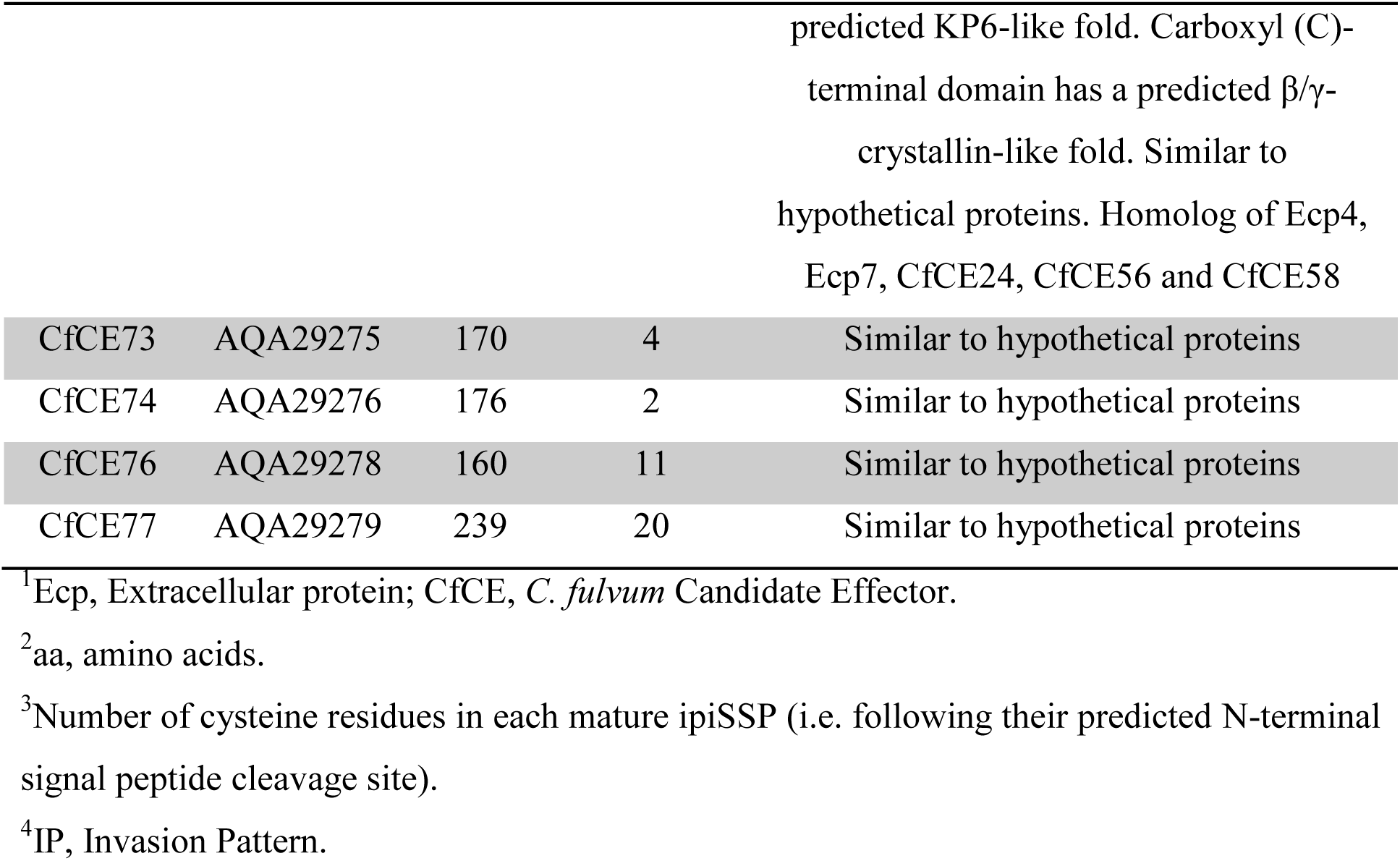
Apoplastic *in planta*-induced small secreted proteins (ipiSSPs) of *Cladosporium fulvum* produced during colonization of susceptible tomato (*Solanum lycopersicum* cv. Heinz-Cf-0).

Amongst the 70 apoplastic ipiSSPs are all *C. fulvum* SSP effectors identified in previous studies (Avr2, Avr4, Avr4E, Avr5, Avr9, Ecp1, Ecp2-1, Ecp4, Ecp5, Ecp6 and Ecp7) (Bolton et al., 2008; Joosten et al., 1994; Laugé et al., 2000; Luderer et al., 2002a; Mesarich et al., 2014; van den Ackerveken et al., 1993; van Kan et al., 1991; Westerink et al., 2004), as well as 32 of 43 (∼74.4%) *C. fulvum* candidate effectors (CfCEs) recently discovered using a combined bioinformatic and transcriptome sequencing approach (Mesarich et al., 2014) (Table S1). The latter includes CfPhiA-1 (CfCE11), a phialide protein previously identified in the IWF sample of a compatible *C. fulvum* (strain IPO 1979)−tomato (*S. lycopersicum* cv. Moneymaker [MM]-Cf-0) interaction at 14 dpi (Bolton et al., 2008).

Strikingly, 62 of the 70 apoplastic ipiSSPs (∼88.6%) are both Cys-rich (≥4 Cys residues) and have an even number of Cys residues (Tables 1 and S1). With the exception of putative propeptide kexin protease cleavage (LXKR) and N-linked glycosylation (NXS/T) sites, no shared motifs were identified between five or more of the 70 ipiSSPs. In total, six ipiSSPs, specifically CfCE16, CfCE20, CfCE33, CfCE40, CfCE66 and CfCE72, possess an LXKR motif (Information S1). In all but one of these ipiSSPs (CfCE72), this motif is located between the predicted signal peptide cleavage site and the first Cys residue (Information S1). A similar motif (LXPR) is located between the predicted signal peptide cleavage site and the first Cys residue of CfCE33 and CfCE67 (Information S1). Twenty-five mature ipiSSPs (∼35.7%) possess one or more NXS/T motifs (Information S1).

Basic local alignment search tool (BLAST) homology searches against publicly available sequence databases at the National Center for Biotechnology Information (NCBI) and the Joint Genome Institute (JGI) revealed that 14 of the 70 apoplastic ipiSSPs are novel (20%), while 47 (∼67.1%) have homology to proteins of unknown function (Tables 1 and S1). The nine remaining ipiSSPs (∼12.9%) have known or predicted functional domains, or have homology to proteins with characterized biological functions. These are: Avr4 (CBM_14 domain; PF01607); Ecp2-1 (Hce2 domain; PF14856); Ecp6 (three LysM domains; PF01476); CfPhiA-1 and CfPhiA-2 (phialide proteins); CfCE55 (class II hydrophobin [Fig. 1]); CfCE60 (GPI-anchored superfamily domain; PF10342); CfCE61 (cerato-platanin protein; PF07249); and CfCE69 (hydrophobic surface-binding protein A [HsbA] domain; PF12296) (Tables 1 and S1). BLASTp homology searches and Cys spacing comparisons also revealed that 23 ipiSSPs are related to each other at the amino acid level. These are: Avr9 and CfCE67; CfCE4 and CfCE16; CfCE5, CfCE25 and CfCE65; CfCE9 and CfCE49; CfCE13 and CfCE63; CfCE14 and CfCE31; CfCE24, CfCE56, CfCE58 and CfCE72 (N-terminal region [NTR; residues 21−113]; CfCE30 and CfCE70 (IgE-binding proteins); CfPhiA-1 and CfPhiA-2; and Ecp4, Ecp7 and CfCE72 (C-terminal region [CTR; residues 158–266]) (Tables 1 and S1).

**Fig. 1.**
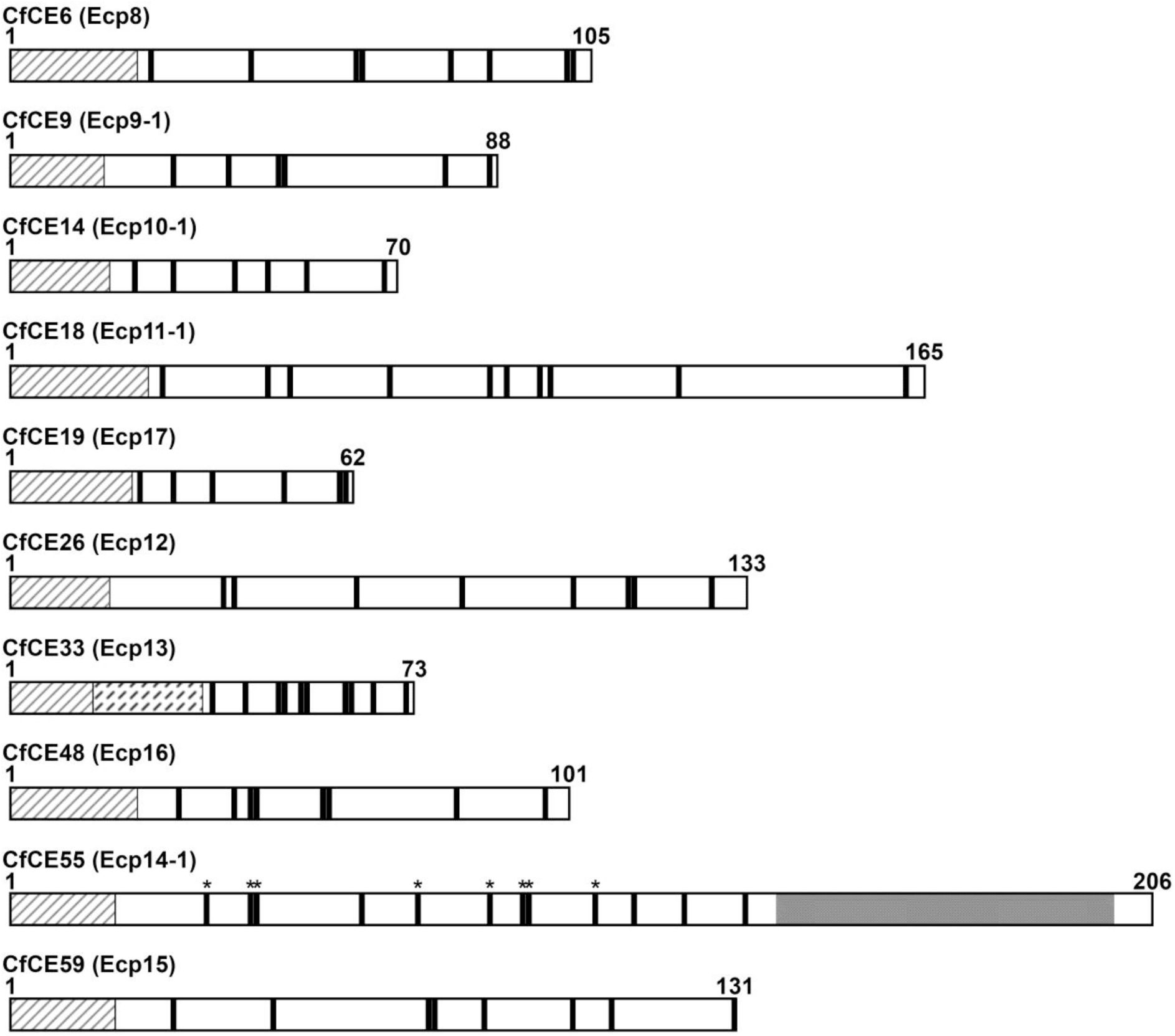
Schematic representation of 10 apoplastic *Cladosporium fulvum* strain 0WU candidate effector (CfCE) proteins that trigger a hypersensitive response (HR) in one or more specific accessions of tomato. All 10 CfCE proteins are small, cysteine-rich, and are predicted to possess an amino (N)-terminal signal peptide for extracellular targeting to the tomato leaf apoplast. The predicted signal peptide of each CfCE protein is shown by black diagonal lines. Cysteine residues are shown by thick vertical bars. Numbers indicate the first and last amino acid residue of each protein. The predicted propeptide domain of CfCE33, ending with a predicted kexin protease cleavage site, is shown by black dashed diagonal lines. A glycine/leucine-rich region present in CfCE55 is shaded grey. Cysteine residues of CfCE55 that are conserved with fungal hydrophobin proteins are shown by asterisks.

As 61 of the 70 apoplastic ipiSSPs (∼87.1%) are novel or have homology to proteins of unknown function, 10 three-dimensional protein structure prediction servers were employed to infer possible structural relationships between these and proteins of characterized tertiary structure and/or function present in the Research Collaboratory for Structural Bioinformatics Protein Data Bank (RCSB PDB). Three ipiSSPs (CfCE5, CfCE25 and CfCE65) were consistently predicted to have structural homology to Alt a 1 (RCSB PDB IDs: 3V0R and 4AUD), an allergen protein with a β−barrel fold (Chruszcz et al., 2012) from the broad host-range Dothideomycete fungal plant pathogen/saprophyte *Alternaria alternata* (Table S2). Four ipiSSPs (Ecp4, Ecp7, CfCE44 and CfCE72 [CTR]) were consistently predicted to have structural homology to proteins with a β/γ-crystallin fold, including the plant antimicrobial protein MiAMP1 from *Macadamia integrifolia* (ID: 1C01) (McManus et al., 1999), and the yeast killer toxin WmKT from *Williopsis mrakii* (ID: 1WKT) (Antuch et al., 1996) (Table S2). A further three ipiSSPs (CfCE24, CfCE56 and CfCE58) were consistently predicted to have structural homology to the α and/or β subunit of KP6 (IDs: 1KP6 and 4GVB), a virus-encoded antifungal killer toxin with an α/β-sandwich fold secreted by the fungal corn smut pathogen *Ustilago maydis* (Allen et al., 2013a; Li et al., 1999) (Table S2). Notably, the NTR of CfCE72 was found to share sequence homology with CfCE24, CfCE56 and CfCE58 (Fig. S1A), suggesting that it too adopts a KP6-like fold. The NTR and CTR of CfCE72 are separated by a putative kexin protease cleavage site (Fig. S1A and Information S1).

Hidden Markov model (HMM)−HMM alignments generated between CfCE5 and Alt a 1, Ecp4 and MiAMP1, as well as CfCE58 and KP6β (i.e. as part of the HHPred server output [Söding et al., 2005]), are shown in Fig. S2. In addition to conserved elements of secondary structure, all three alignments revealed conserved Cys residues. For CfCE5 and Alt a 1, two conserved Cys residues at positions 50 and 65 (mature proteins), which are also present in CfCE25 and CfCE65, were identified (Figs S1B and S2A). In Alt a 1, these Cys residues are known to form an intramolecular disulphide bond (Chruszcz et al., 2012). Inspection of the predicted CfCE5 tertiary structure, which was modelled using Alt a 1 as a template in HHpred (MODELLER) (Söding et al., 2005; Webb and Sali, 2002) and RaptorX (Källberg et al., 2012), suggests that the conserved Cys50/Cys65 pair forms an intramolecular disulphide bond (Fig. S3A). Furthermore, the predicted structure suggests that the two remaining Cys residues, Cys24 and Cys29, which are absent from Alt a 1 (Fig. S2A), may also form an intramolecular disulphide bond, given that they are located in close proximity to each other (Fig. S3A). This bond, however, would be located in a different location to the second intramolecular disulphide bond of Alt a 1 (Cys104−Cys116) (Fig. S3A) (Chruszcz et al., 2012).

Five of the six Cys residues present in Ecp4 and MiAMP1 were found to be conserved (Fig. S2B). In MiAMP1, all six Cys residues are known to form intramolecular disulphide bonds (Cys11−Cys65, Cys21–Cys76 and Cys23–Cys49) (McManus et al., 1999). Inspection of the predicted Ecp4 structure, which was modelled using MiAMP1 as a template, suggests that two of the conserved Cys pairs, Cys16/Cys84 and Cys35/Cys67, form intramolecular disulphide bonds (Fig. S3B). Although not conserved, the sixth Cys residue in Ecp4, Cys57, still appears to be located in a favourable position for disulphide bond formation with Cys99 (Fig. S3B). All six Cys residues in Ecp4 are conserved across Ecp7 and CfCE72 (CTR), although the latter has an additional pair of Cys residues (Fig. S1C).

For CfCE58 and KP6β, six conserved Cys residues, which are also present in CfCE24 and CfCE56, were identified (Figs S1A and S2C). In KP6β, these six Cys residues are known to form three intramolecular disulphide bonds (Cys9−Cys74, Cys11−Cys64 and Cys29−Cys46) (Allen et al., 2013a). The predicted CfCE58 structure, which was modelled using KP6β as a template, suggests that the three conserved Cys pairs (Cys7/Cys76, Cys9/Cys66 and Cys26/Cys47) form intramolecular disulphide bonds (Fig. S3C). Both CfCE56 and CfCE58 possess an additional set of Cys residues (Cys1 and Cys60) (Fig. S1A). Cys1 of CfCE58 is located at the extreme N-terminus, which, if flexible, would be expected to make contact with Cys60 located at the base of one of the predicted α-helices (Fig. S3C).

### Most apoplastic ipiSSPs of *C. fulvum* lack an ortholog in *Dothistroma septosporum*

Of the fungi for which a genome sequence is so far available, *D. septosporum* is the most closely related to *C. fulvum* (de Wit et al., 2012). Reciprocal BLASTp and tBLASTn searches were used to determine whether the predicted *D. septosporum* protein catalogue and genome (de Wit et al., 2012) carry homologs of the 70 *C. fulvum* apoplastic ipiSSPs and their encoding genes, respectively. For 43 of the 70 ipiSSPs, no homologs were identified (Table S1). A further four showed limited homology to *D. septosporum* genes, while five others had homology to pseudogenes (Table S1). The remaining 18 ipiSSPs had likely orthologs in *D. septosporum*. However, of these, only 11 were up-regulated during infection of pine (Table S1) (Bradshaw et al., 2016). More specifically, these are the likely orthologs of Ecp2-1, Ecp6, CfCE33, the three Alt a 1 allergen-like proteins (CfCE5, CfCE25 and CfCE65), CfCE16, the cerato-platanin (CfCE61), the phialide protein CfPhiA-2 (CfCE53), CfCE74 and CfCE77 (Table S1). Genes encoding SSPs with a potential β/γ−crystallin or KP6-like fold were absent, pseudogenized, or not expressed during colonization of pine (Table S1).

### Nine apoplastic ipiSSPs of *C. fulvum* trigger an HR in specific accessions of tomato

To identify new sources of resistance against *C. fulvum*, wild accessions of tomato were screened for their ability to recognize apoplastic ipiSSPs using the PVX-based transient expression system (Hammond-Kosack et al., 1995; Takken et al., 2000). In this experiment, recombinant viruses were delivered through agroinfection for local (toothpick wounding) or systemic (cotyledon infiltration) expression of ipiSSPs in tomato, with the pathogenesis-related 1A (PR1A) signal peptide of tobacco (*Nicotiana tabacum*) used to direct secretion of these proteins into the tomato leaf apoplast. Plants that showed a chlorotic or necrotic HR were deemed to have recognized an ipiSSP as an IP.

As a starting point, 25 predominantly wild accessions of tomato (Table S3) were screened for their ability to recognize Ecp7 and/or one or more of 40 CfCEs (Table S1) using the PVX agroinfection method based on toothpick wounding (Luderer et al., 2002a; Takken et al., 2000). This set of 40 CfCEs primarily comprises those with the highest level of expression *in planta*, as based on pre-existing RNA-Seq data shown in Table S1. A fully expanded leaf from 1−3 representative plants of each accession was inoculated via toothpick wounding on each side of the main vein, and the presence or absence of an HR was scored at 10 dpi. At the same time, *S. lycopersicum* cv. MM-Cf-0 (no *Cf* immune receptors; Tigchelaar, 1984) was screened to determine whether Ecp7 or any of the CfCEs trigger a non-specific HR. Likewise, accessions carrying only the *Cf-1, Cf-3, Cf-6, Cf-9B, Cf-11* or *Cf-Ecp3* immune receptor gene (Table S3) were screened to determine whether Ecp7 or any of the CfCEs represent one of the yet unknown IP effectors Avr1, Avr3, Avr6, Avr9B, Avr11 or Ecp3. As positive controls, *S. lycopersicum* cv. MM-Cf-5, which carries only the *Cf-5* immune receptor (Tigchelaar, 1984), as well as the landrace accession CGN 18399 (*S. lycopersicum* var. *cerasiforme*), from which the *Cf-5* gene was originally identified (Kerr et al., 1971), were screened for their ability to recognise the IP effector Avr5 (Mesarich et al., 2014). Empty vector was used as a negative control to confirm that PVX alone does not trigger a non-specific HR. For the purpose of this experiment, recognition of Ecp7 or a CfCE was deemed to have occurred if an HR was triggered at one or both of the toothpick wounding sites on a given tomato leaf.

As expected, the empty vector (negative control) failed to trigger an HR in any tomato accession tested, while Avr5 (positive control) was recognized by only MM-Cf-5 and CGN 18399 (Fig. S4), indicating that the PVX agroinfection method is functional, and that no other accessions carry the *Cf-5* immune receptor gene. Ten of the 40 CfCEs (CfCE6, CfCE9, CfCE14, CfCE18, CfCE19, CfCE26, CfCE33, CfCE48, CfCE55 and CfCE59) were recognized by one to eight predominantly wild accessions of tomato, with HRs ranging from weak chlorosis to strong necrosis (Fig. S4). Furthermore, 15 of the 25 accessions recognized between one and four of the 10 CfCEs (Fig. S4). Importantly, none of the 10 CfCEs triggered an HR in MM-Cf-0, suggesting that the observed responses were specific to the accessions tested (Fig. S4). None of the accessions carrying the *Cf-1, Cf-3, Cf-6, Cf-9B, Cf-11* or *Cf-Ecp3* immune receptor gene recognized Ecp7 or any of the CfCEs, indicating that these ipiSSPs do not represent the IP effectors Avr1, Avr3, Avr6, Avr9B, Avr11 or Ecp3. A schematic of the 10 HR-eliciting CfCEs is shown in Fig. 1.

To further confirm recognition of the 10 CfCEs, each was screened for its ability to trigger a systemic HR in the same responding tomato accessions using the PVX agroinfection method based on cotyledon infiltration (Mesarich et al., 2014). Here, both cotyledons of five independent plants were infiltrated, and the presence or absence of an HR was scored at 21 dpi. Consistent with the agroinfection assay based on toothpick wounding, the empty vector (negative control) did not trigger an HR in any accession tested (Figs 2 and S5). Similarly, none of the CfCEs triggered an HR in MM-Cf-0 (Fig. S6). For CfCE6, CfCE26, CfCE48 and CfCE55, recognition could be confirmed across all responding accessions identified in the toothpick wounding agroinfection assay (Figs 2 and S7−S8). Recognition could also be confirmed across most, but not all, previously identified accessions for CfCE9, CfCE14, CfCE18, CfCE33 and CfCE59 (Figs 2 and S9–S13). Indeed, CfCE9, CfCE14, CfCE18 and CfCE33 only failed to trigger an HR in accessions CGN 15392 (*Solanum arcanum*) (Fig. S9), CGN 14356 (*Solanum peruvianum*) (Fig. S10), CGN 14357 (*Solanum corneliomuelleri*) (Fig. S11) and CGN 14353 (*S. pimpinellifolium*) (Fig. S12), respectively, while CfCE59 only failed to trigger an HR in CGN 14353 and CGN 24034 (*S. pimpinellifolium*) (Fig. S13). In some cases, the recognition of a CfCE could not be observed across all five plants of a given accession representing *S. corneliomuelleri* (CGN 14357 and CGN 15793), *S. peruvianum* (CGN 14355, CGN 14356 and CGN 24192) and *S. pimpinellifolium* (CGN 15946) (Figs 2, S7, S9, S11 and S13). In all responding accessions, the systemic HR involved weak to strong necrosis, and was typically associated with moderate to severe stunting (Figs 2, S7−S11 and S^13^–S14). The recognition of only one CfCE, CfCE19, could not be confirmed (CGN 24034; Fig. S15).

**Fig. 2.**
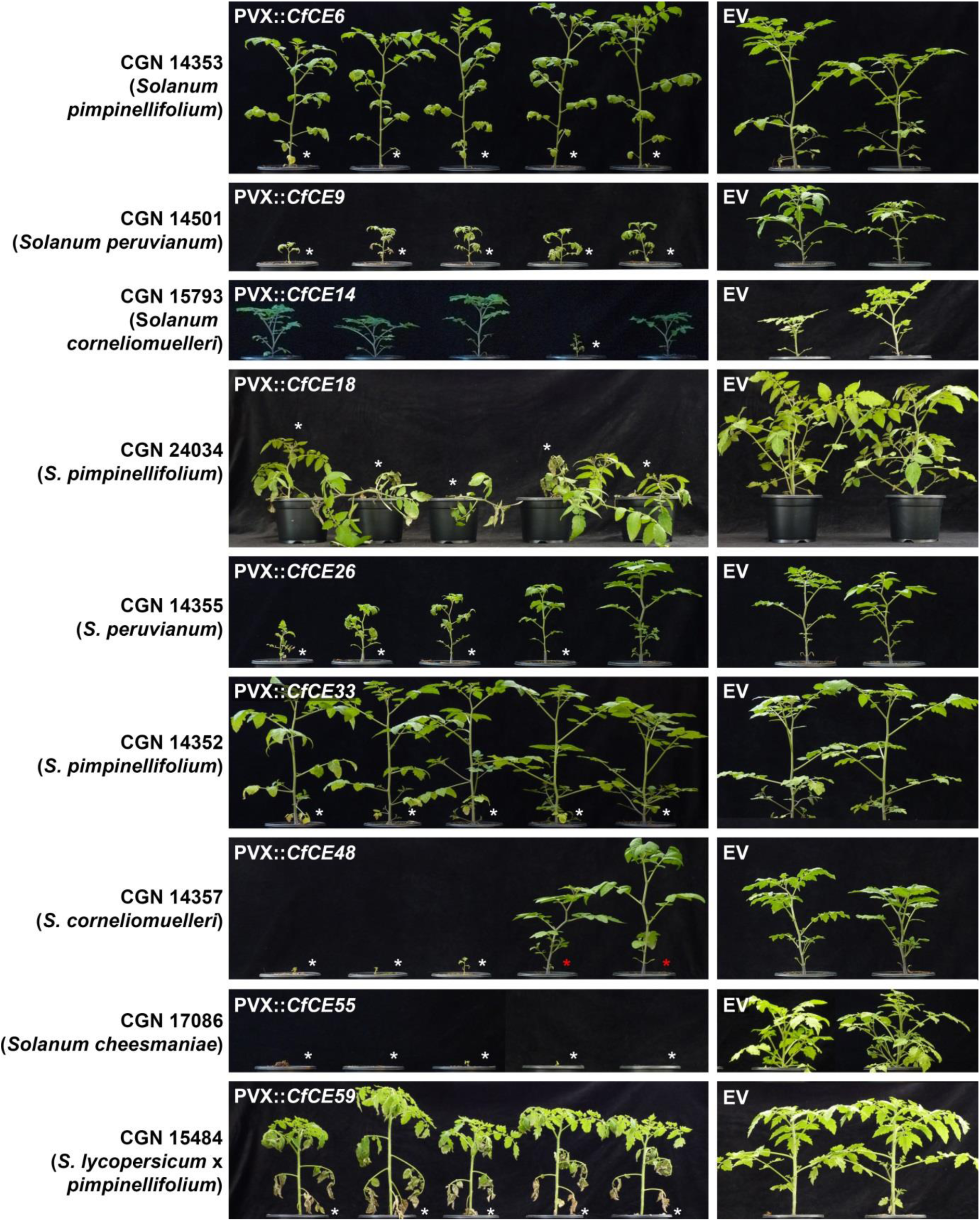
Nine *Cladosporium fulvum* candidate effectors (CfCEs) of strain 0WU trigger a systemic hypersensitive response (HR) in one or more specific accessions of tomato. Selected examples are shown. CfCEs were systemically produced in five representatives of each tomato accession (left) using the *Potato virus X* (PVX) transient expression system. Recombinant PVX was delivered by *Agrobacterium tumefaciens* (agroinfection) through cotyledon infiltration at 10 d post-seed germination. Two representatives of each tomato On were inoculated with PVX alone (pSfinx empty vector; EV) (right). Plants ing a systemic chlorotic or necrotic HR are shown by white asterisks. Plants without s mosaic symptoms (i.e. not infected with PVX) are shown by red asterisks. raphs were taken at 21 d post-infiltration.

### Tomato accessions that recognize apoplastic ipiSSPs are resistant to *C. fulvum*

To determine whether the accessions of tomato that recognize apoplastic ipiSSPs are resistant to *C. fulvum*, each, along with *S. lycopersicum* cv. MM-Cf-0, was inoculated with strain 2.4.5.9.11 IPO of this fungus, and symptoms were inspected on leaves from three independent plants at 14 dpi. Strain 2.4.5.9.11 IPO carries genes corresponding to all nine HR-eliciting CfCEs (see below), but lacks a functional copy of the previously cloned *Avr2, Avr4, Avr4E, Avr5* and *Avr9 IP* effector genes (Mesarich et al., 2014; Stergiopoulos et al., 2007). As expected, *S. lycopersicum* cv. MM-Cf-0 was susceptible to 2.4.5.9.11 IPO (Fig. S16). In contrast, all other tomato accessions tested were resistant to this strain (Fig. S16). For accessions CGN 14474 (*S. lycopersicum*) and CGN 15820 (*S. lycopersicum* × *cheesmaniae*), this resistance was observed across only two of the three independent plants (Fig. S16). While resistant to *C. fulvum*, we cannot exclude the possibility that the set of resistant tomato accessions carries one or more of, for example, the *Cf* immune receptor genes *Cf-1, Cf-3, Cf-6, Cf-9B, Cf-Ecp1, Cf-Ecp2-1, Cf-Ecp3, Cf-Ecp4, Cf-Ecp5* and *Cf-Ecp6*.

As CfCE6, CfCE9, CfCE14, CfCE18, CfCE26, CfCE33, CfCE55, CfCE59 and CfCE48 are present in IWF samples from compatible *C. fulvum*–tomato interactions, and because these proteins triggered an HR using both PVX agroinfection methods, only these apoplastic ipiSSPs were pursued further. From this point forward, CfCE6, CfCE9, CfCE14, CfCE18, CfCE26, CfCE33, CfCE55, CfCE59 and CfCE48 will be referred to as Ecp9-1, Ecp10-1, Ecp11-1, Ecp12, Ecp13, Ecp14-1, Ecp15 and Ecp16, respectively.

### Seven HR-eliciting Ecps have one or more homologs in other fungal species, while three HR-eliciting Ecps have one or more paralogs in *C. fulvum*

To identify homologs of the HR-eliciting Ecps in other fungi, each was screened against the publicly available protein sequence databases at NCBI and JGI using BLASTp. Additionally, in those cases where no protein homolog could be identified, Ecps were screened against the collection of fungal genome sequences present at JGI using tBLASTn (i.e. to identify homologs without a gene prediction). With the exception of Ecp8 and Ecp16, homologs of all HR-eliciting Ecps were identified in other fungal species. For Ecp9-1, homologs were identified in the Dothideomycetes *Pseudocercospora fijiensis* (black sigatoka disease of banana), *Septoria musiva* and *Septoria populicola* (leaf spot and canker diseases of poplar), *Teratosphaeria nubilosa* (leaf spot of *Eucalyptus* spp.) and *Zasmidium cellare* (saprobic wine cellar fungus), as well as eight Sordariomycete species (Fig. S17). Eight paralogs of Ecp9-1 were found to be encoded by the genome of *C. fulvum* strain 0WU (Ecp9-2−Ecp9-9) (Fig. S18A), with one clear pseudogene also identified (*Ecp9-10*; *result not shown*). A similar expansion was found in the Sordariomycete *Claviceps purpurea* (ergot disease of cereals) (Fig. S17).

Homologs of Ecp10-1 were identified in the Dothideomycetes *Pseudocercospora eumusae* and *Pseudocercospora musae* (eumusae leaf spot and yellow sigatoka disease of banana, respectively), *A. alternata, S. musiva, S. populicola, T. nubilosa* and *Z. cellare*, as well as *Zymoseptoria ardabiliae, Zymoseptoria pseudotritici* and *Zymoseptoria tritici* (leaf blotch diseases of grasses), *Venturia inaequalis* and *Venturia pirina* (apple and pear scab disease, respectively), *Clathrospora elynae* (found growing on curved sedge), *Cochliobolus sativus* and *Cochliobolus victoriae* (cereal pathogens), *Pyrenophora teres* f. *teres* (net blotch disease of barley), *Pyrenophora tritici-repentis* (tan spot disease of wheat) and *Setosphaeria turcica* (northern corn leaf blight disease) (Fig. S19 and Information S2). Homologs of Ecp10-1 were also identified in several Sordariomycete fungi (Fig. S19 and Information S2). Interestingly, Ecp10-1 homologs were found to be massively expanded in *V. inaequalis* and *V. pirina* (Information S2), which is not uncommon for effector candidates from these fungi (Deng et al., 2017). Smaller expansions were also identified in other fungal plant pathogens (Information S2). Two paralogs of Ecp10-1 (Ecp10-2 and Ecp10-3) were found to be encoded by the genome of *C. fulvum* strain 0WU (Fig. S18B).

Homologs of the remaining Ecps were only identified in Dothideomycete fungi. Ecp11-1 was found to have homology to AvrLm3 and AvrLmJ1, two avirulence effector proteins from *Leptosphaeria maculans* (blackleg disease of Brassica species) (Plissonneau et al., 2016; van de Wouw et al., 2014), as well as two proteins from *Z. ardabiliae* (Figs 3 and S20). A single pseudogene of *Ecp11-1* (*Ecp11-2*) was also identified in the genome of *C. fulvum* strain 0WU (*result not shown*). Ecp12 was found to have multiple homologs in *S. musiva* and *S. populicola*, with the homologous Cys-rich domain occurring once, or as two or three tandem repeats (Fig. S21), as has been found for several other effectors from plant-associated organisms (Mesarich et al., 2015). Homologs of Ecp13 were identified in *D. septosporum, P. fijiensis, S. musiva* and *Cercospora zeae-maydis* (grey leaf spot disease of maize) (Fig. S22), while homologs of Ecp14-1 were found in *C. zeae-maydis, D. septosporum, P. eumusae P. fijiensis, P. musae, S. musiva, S. populicola, T. nubilosa, Trypethelium eluteriae* (lichen-forming fungus), *Z. ardabiliae, Zymoseptoria brevis* (leaf blotch disease of barley), *Z. pseudotritici, Z. tritici* and *Z. cellare*, with most, including *C. fulvum*, possessing a paralog (Figs S18C and S23). A single pseudogene of *Ecp14-1* (*Ecp14-3*) was identified in the genome of *C. fulvum* strain 0WU (*result not shown*). For Ecp15, homologs were found in *P. fijiensis, P. musae* and *Z. ardabiliae* (Fig. S24).

**Fig. 3.**
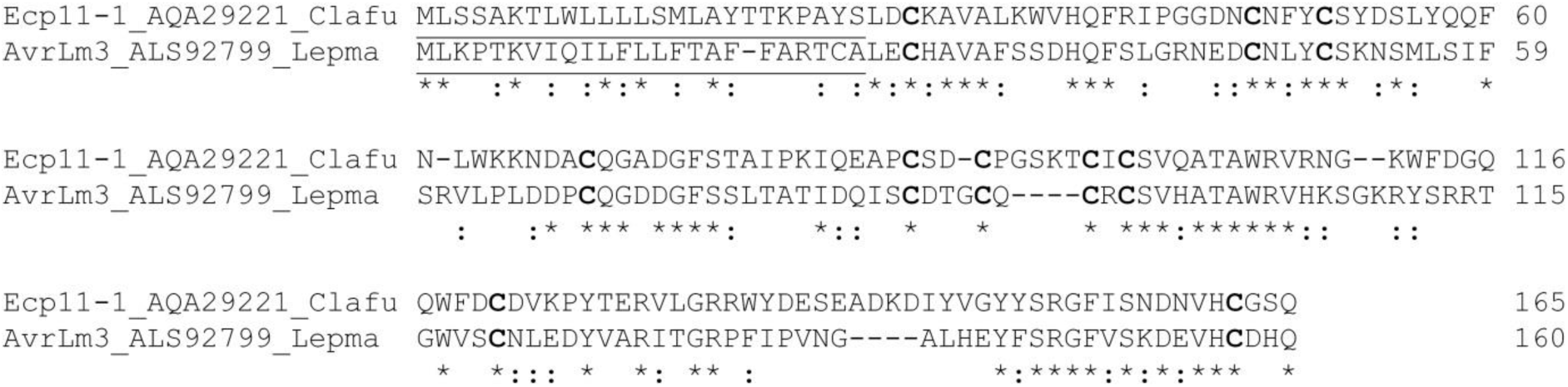
Ecp11-1 of *Cladosporium fulvum* is a homolog of AvrLm3 from *Leptosphaeria maculans*. Conserved (*) and physicochemically similar (:) amino acid residues shared between Ecp11-1 and AvrLm3 are shown below the alignment. Cysteine residues are highlighted in bold. The predicted amino (N)-terminal signal peptide sequence of Ecp11-1 and AvrLm3 is underlined.

### Genes encoding HR-eliciting Ecps are induced *in planta*

RNA-Seq fragments per kilobase (kb) of exon per million fragments mapped (FPKM) values suggested that all genes encoding an HR-eliciting Ecp of *C. fulvum*, like those encoding all previously identified IP effectors of this fungus (Mesarich et al., 2014), are induced during infection of susceptible tomato, when compared to expression during growth *in vitro* in PDB or Gamborg B5 liquid media (Table S1). To confirm this expression profile, a reverse-transcription quantitative real-time polymerase chain reaction (RT-qrtPCR) experiment was performed. Indeed, all genes encoding an HR-eliciting Ecp were found to be induced during infection of susceptible tomato, when compared to expression during growth *in vitro* in PDB or Gamborg B5 liquid media (Fig. 4).

**Fig. 4.**
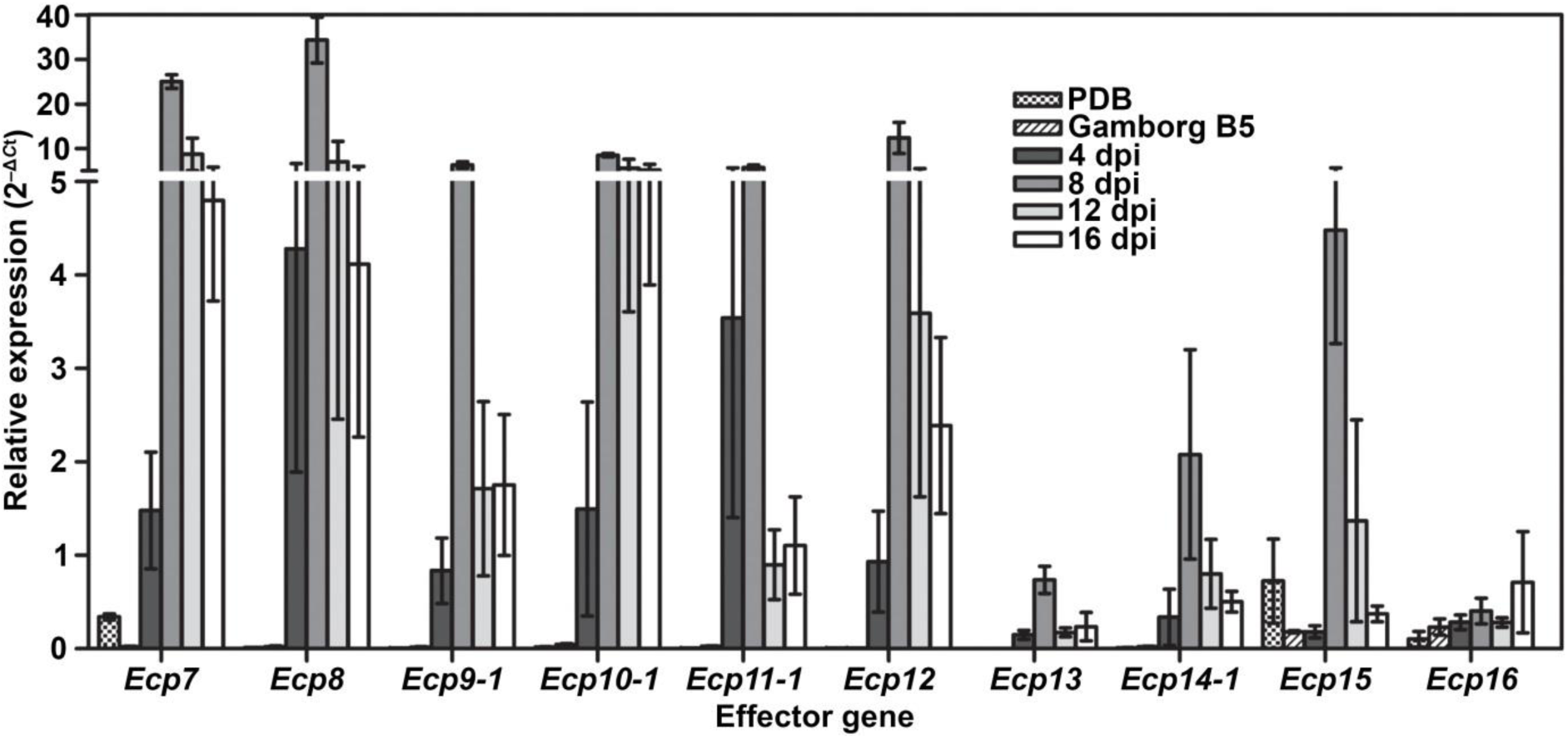
Genes encoding a hypersensitive response (HR)-eliciting extracellular protein (Ecp) from *Cladosporium fulvum* strain 0WU are induced *in planta*. Expression was monitored by a reverse-transcription–quantitative real-time polymerase chain reaction (RT-qrtPCR) experiment *in planta* during a compatible *C. fulvum* strain 0WU–*Solanum lycopersicum* cv. Heinz Cf-0 interaction at 4, 8, 12 and 16 d post-inoculation (dpi), as well as during growth of *C. fulvum* strain 0WU *in vitro* in potato-dextrose broth (PDB) and Gamborg B5 liquid media at 4 dpi. The *C. fulvum actin* gene was targeted for normalisation of expression, which was calculated using the 2−Δ*C*t method. Error bars represent the standard deviation of three biological replicates.

### Most genes encoding an HR-eliciting Ecp are associated with repetitive elements

It is common for *C. fulvum* effector genes to be flanked by a mosaic of repetitive elements in the genome of strain 0WU (de Wit et al., 2012; Mesarich et al., 2014). It has been proposed that these elements may assist in the deletion of *IP* effector genes following *Cf* immune receptor-imposed selection pressure (Mesarich et al., 2014). To determine whether repetitive elements also flank genes encoding the HR-eliciting Ecps, the genome scaffolds harbouring each of these genes was screened for repetitive sequence across the *C. fulvum* 0WU genome using BLASTn. Six of the nine *Ecp* genes (*Ecp8, Ecp9-1, Ecp10-1, Ecp11-1, Ecp12* and *Ecp15*) were found to be associated with repetitive elements at both their 5′ and 3′ flanks (Fig. S25). Furthermore, the same six genes were found to reside on small genome scaffolds of less than 35 kb in length (Table S4). The latter suggests that the scaffolds harbouring these genes are surrounded by even larger flanking repetitive elements, with these elements anticipated to have hampered a larger scaffold assembly (Wit et al., 2012). The 5′ end of *Ecp16* is closely associated with repetitive elements, and is present at the 5′ end of an ∼55-kb scaffold (Fig. S25). Likewise, *Ecp13* is located at the 3′ end of an ∼57-kb scaffold, suggesting the presence of 3′ repeats (Fig. S25). In contrast to the *Ecp* genes mentioned above, *Ecp14-1* is not surrounded by repetitive elements (Fig. S25).

### Genes encoding an HR-eliciting Ecp exhibit limited allelic variation between strains

It is common for genes encoding *C. fulvum* IP effectors to exhibit allelic variation between strains, which is often brought about by selection pressure to avoid recognition by corresponding Cf immune receptors (Iida et al., 2015; Joosten et al., 1994; Luderer et al., 2002a; Mesarich et al., 2014; Westerink et al., 2004). To assess the level of allelic variation across genes encoding the HR-eliciting Ecps, each was amplified by PCR from 10 different *C. fulvum* strains (Table S5), sequenced, and compared to the corresponding sequence from strain 0WU. All nine *Ecp* genes could be amplified by PCR from genomic DNA samples representing the 10 *C. fulvum* strains. Of the nine genes, four, namely *Ecp9-1, Ecp10-1, Ecp13* and *Ecp15*, exhibited no allelic variation between strains. For *Ecp8* and *Ecp16*, allelic variation was observed; however, this variation did not result in a change of amino acid sequence. More specifically, in six strains (2.4, 2.4.5, 2.5, 2.9, 4 and 7320), *Ecp8* had a single synonymous CCC→CCT substitution at position 153, while in four strains (2.4, 2.4.5, 2.5 and 4), *Ecp16* had a trinucleotide insertion (CTT) at position 234 in an intron (Fig. 5). For each of the remaining three genes, a single non-synonymous substitution was identified: a TTT→GTT (Phe119Val) change at position 355 in *Ecp11-1* of strain 2.9; a GGG→AGG (Gly124Arg) change at position 484 in *Ecp12* of strains 2.9 and 7320; and an AAG→GAG (Lys148Glu) change at position 501 in *Ecp14-1* of strains 2.4, 2.4.5, 2.4.5.9.11 IPO, 2.4.9.11, 2.5, 2.9 and 4 (Fig. 5). A G→T mutation at position 386 of the *Ecp14-1* intron in strains 2.4, 2.4.5, 2.4.5.9.11 IPO, 2.4.9.11, 2.5 and 4, as well as a synonymous GGG→GGA substitution at position 452 in *Ecp14-1* of strains 2.4, 2.4.5, 2.4.5.9.11 IPO, 2.4.9.11, 2.5, 2.9 and 4, were also identified (Fig. 5). It is not yet known whether the non-synonymous substitutions identified in *Ecp11-1, Ecp12* and *Ecp14-1* allow *C. fulvum* to overcome resistance mediated by the putative *Cf-Ecp11-1, Cf-Ecp12* and *Cf-Ecp14-1* immune receptor genes, respectively.

**Fig. 5.**
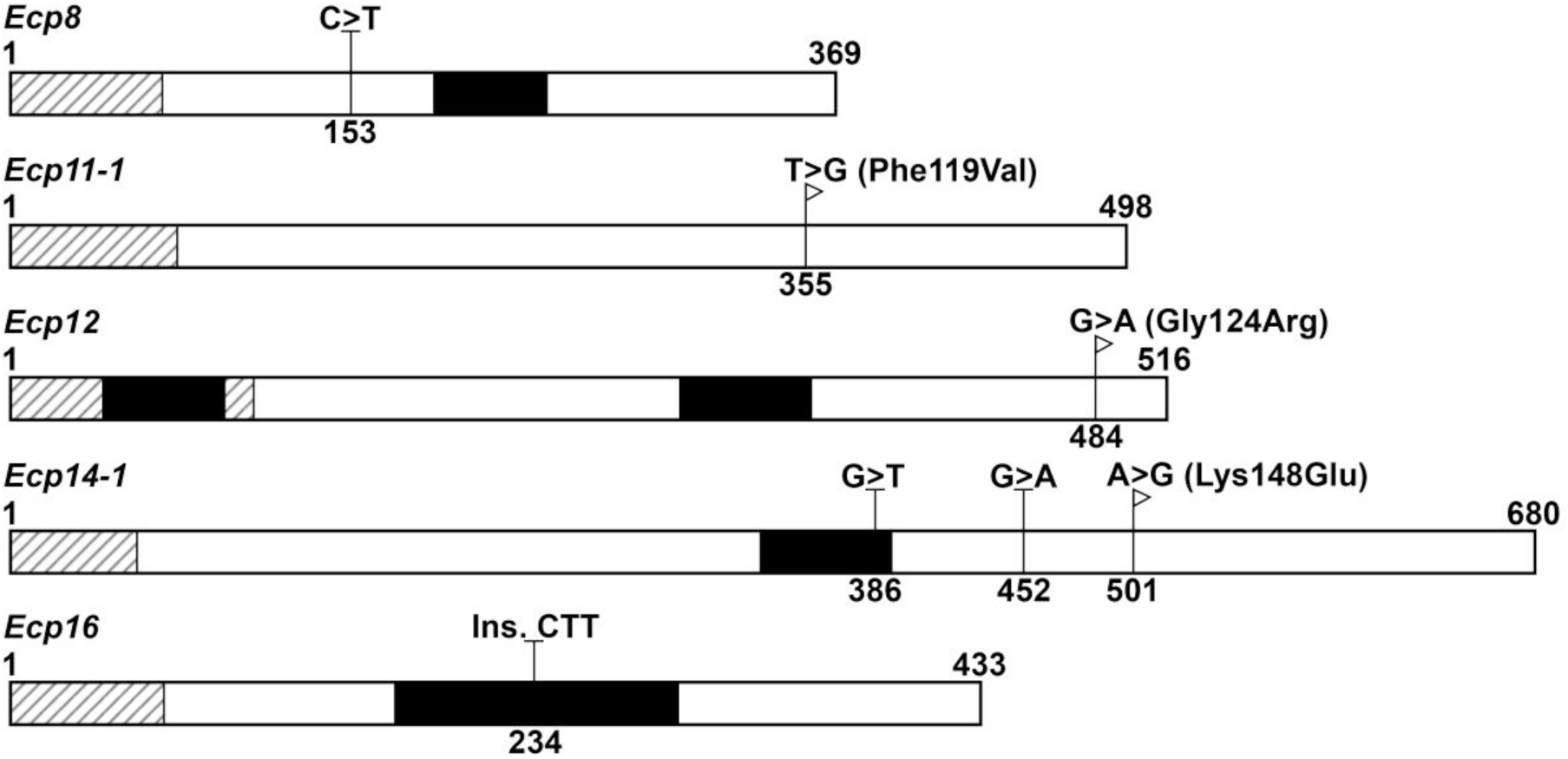
Genes encoding a hypersensitive response (HR)-eliciting extracellular protein (Ecp) exhibit limited allelic variation between strains of *Cladosporium fulvum*. Allelic variation was assessed across 10 distinct strains of *C. fulvum*, and was compared to strain 0WU. Open reading frames (encoding each mature protein) and introns are shown as white and black boxes, respectively. Regions of each *Ecp* gene predicted to encode an amino (N)-terminal signal peptide sequence are shown by black diagonal lines. DNA modifications leading to non-synonymous amino acid substitutions are shown by white flags. DNA modifications leading to synonymous amino acid mutations or changes to intronic sequences are shown by Ts. Numbers above each schematic represent the first and last nucleotide of each gene (i.e. of the ATG to STOP codons, respectively). Numbers on the bottom of each schematic represent the location of each DNA modification.

## DISCUSSION

Leaf mould disease of tomato, caused by the fungal pathogen *C. fulvum*, is a re-emerging problem worldwide. This re-emergence is due to intensive year-round cultivation of resistant tomato cultivars, which have selected for natural strains of this fungus capable of overcoming, for example, one or more of all cloned *Cf* immune receptor genes (Hubbeling, 1978; Iida et al., 2015; Laterrot, 1986; Li et al., 2015). To combat these strains, new *Cf* immune receptor genes need to be identified. Wild tomato is a rich source of resistance against *C. fulvum* (Kruijt et al., 2005; Laugé et al., 1998, 2000; van der Hoorn et al., 2001b). In this study, an effectoromics approach (Domazakis et al., 2017; Du and Vleeshouwers, 2014) based on apoplastic ipiSSPs of *C. fulvum* was used to identify wild accessions of tomato carrying new *Cf* immune receptor genes.

As a starting point for this approach, proteomics and transcriptome sequencing were used to identify fungal SSPs most relevant to the *C. fulvum*–tomato interaction. Altogether, 70 apoplastic ipiSSPs, made up of all 11 characterized SSP effectors of this fungus (Bolton et al., 2008; Joosten et al., 1994; Laugé et al., 2000; Luderer et al., 2002a; Mesarich et al., 2014; van den Ackerveken et al., 1993; van Kan et al., 1991; Westerink et al., 2004), as well as 32 previously described (Mesarich et al., 2014) and 27 new CfCEs, were identified in IWF samples from compatible *C. fulvum*−*S. lycopersicum* cv. H-Cf-0 interactions. Strikingly, all but eight of these ipiSSPs are Cys-rich and possess an even number of Cys residues. Consistent with that shown for Avr4, Avr9, Ecp1, Ecp2-1, Ecp5 and Ecp6, it is likely that many of these Cys residues form intramolecular disulphide bonds required for stability and function in the protease-rich leaf apoplast of tomato (Joosten et al., 1997; Luderer et al., 2002b; Sánchez-Vallet et al., 2013; van den Burg et al., 2003; van den Hooven et al., 2001).

Following signal peptide cleavage, several of the ipiSSPs likely undergo further post-translational processing in the ER–Golgi secretory pathway. Twenty-five ipiSSPs possess one or more NXS/T motifs following their predicted signal peptide cleavage site, suggesting that they undergo N-linked glycosylation. This glycosylation may be required for ipiSSP folding, structure, stability, solubility, oligomerization, or function (Helenius and Aebi, 2001). A further six ipiSSPs possess a putative N-terminal kexin protease cleavage site (LXK/PR motif), suggesting that they have a propeptide domain. It is possible that these ipiSSPs are synthesized as inactive precursors, and that, for biological activity, their propeptide domain must be removed by a kexin protease (Rockwell et al., 2002).

BLAST homology searches revealed that, in addition to Avr4 (single CBM_14 domain; PF01607) (van den Burg et al., 2003), Ecp2-1 (single Hce2 domain; PF14856) (Stergiopoulos et al., 2012), Ecp6 (three LysM domains; PF01476) (Bolton et al., 2008) and CfPhiA-1 (phialide protein) (Bolton et al., 2008), five other ipiSSPs, specifically CfPhiA-2, CfCE60, CfCE61, CfCE69 and Ecp14-1, possess a known functional domain or have homology to proteins with a characterized biological function. Of these, CfPhiA-2 has homology to CfPhiA-1 and other phialide proteins from Ascomycete fungi. To date, the best characterized of these homologs is PhiA from *Aspergillus nidulans*, which localizes to the cell wall of phialides and conidia (Melin et al., 2003). PhiA plays an essential role in the development of phialides, which are sporogenous cells that produce and release conidia through a specialized apical budding process (Melin et al., 2003).

CfCE60 has a GPI-anchored superfamily domain (PF10342), but is not predicted to possess a GPI anchor modification site. Little functional information is available for secreted proteins with this domain. However, in the Basidiomycete fungus *Lentinula edodes* (shiitake mushroom), the PF10342 domain-containing protein Le.DRMIP, which also possesses a mitochondrial targeting signal peptide and transmembrane domain, interacts with the developmentally regulated MAP kinase Le.MAPK. Both proteins have been proposed to play a role in cell differentiation during fruiting body development (Szeto et al., 2007).

CfCE61 is a member of the cerato-platanins (PF07249), a class of proteins ubiquitous to filamentous fungi that adopts a double Ψβ-barrel fold similar to domain one of expansins (Chen et al., 2013; de Oliveira et al., 2011). Cerato-platanins are predominantly secreted, although several also localize to the cell wall of ascospores, conidia and hyphae (e.g. Boddi et al., 2004; Pazzagli et al., 1999). Cerato-platanins are postulated to carry out multiple biological functions related to fungal growth and development, as well as to plant−fungus interactions. Notably, cerato-platanins bind chitin, but not cellulose (Baccelli et al., 2014; de O. Barsottini et al., 2013; Frischmann et al., 2013), yet several members have expansin-like activity *in vitro*, loosening cellulosic materials (Baccelli et al., 2014; de O. Barsottini et al., 2013). It has thus been hypothesized that cerato-platanins may function as expansins required for fungal cell wall remodelling and enlargement, possibly by disrupting non-covalent interactions between β−glucan or chitin molecules (de Oliveira et al., 2011). Epl1, a surface-active cerato-platanin from the biocontrol agent *Trichoderma atroviride*, self-assembles at the air/water interface, forming protein films that increase the polarity of solutions and surfaces (Frischmann et al., 2013). This suggests an additional role for cerato-platanins in increasing the wettability of hyphae, enabling them to grow in aqueous environments, or in protecting them from desiccation (Frischmann et al., 2013).

Deletion of the gene encoding MSP1, a cerato-platanin from the rice blast pathogen *Magnaporthe oryzae*, resulted in reduced virulence *in planta*, suggesting that certain members of this protein class function as effectors (Jeong et al., 2007). In line with this, preliminary studies have suggested that MpCP5, a cerato-platanin from *Moniliophthora perniciosa* (witches’ broom disease of cocoa) may, like Ecp6, perturb chitin-triggered immunity (de O. Barsottini et al., 2013), while cerato-platanins from *Fusarium graminearum* (cereal head blight disease) may, like Avr4, protect fungal cell wall polysaccharides from enzymatic digestion by chitinases and β-1,3-glucanases (Quarantin et al., 2016). Some cerato-platanins are also well-known IPs that trigger a non-specific HR upon recognition by corresponding host immune receptors (e.g. Frías et al., 2011, 2014). This, however, does not appear to be the case for CfCE61, which failed to trigger an HR in tomato.

CfCE69 contains an HsbA domain (PF12296), which was originally identified in the HsbA protein from *Aspergillus oryzae* (Ohtaki et al., 2006), a filamentous fungus commonly used in the fermentation industry. In culture, HsbA is secreted in the presence of the hydrophobic polymer polybutylene succinate-*co*-adipate (PBSA). HsbA binds PBSA, and in doing so, recruits CutL1, a polyesterase/cutinase, for its degradation (Ohtaki et al., 2006).

Ecp14-1 is a member of the hydrophobins, a fungal-specific class of surface-active proteins (Wessels, 1994). With the exception of eight conserved Cys residues, which form four intramolecular disulphide bonds, hydrophobins share limited sequence similarity (Wessels, 1994). Ecp14-1 is the twelfth hydrophobin, and sixth class II hydrophobin, to be identified from *C. fulvum* (de Wit et al., 2012; Nielsen et al., 2001; Segers et al., 1999; Spanu, 1997). It is also the first hydrophobin to be identified from this fungus that is exclusively expressed *in planta* (Fig. 4). Hydrophobins are initially secreted in a soluble form, but then spontaneously localize to hydrophilic:hydrophobic interfaces, where they assemble into insoluble, amphipathic layers (Sunde et al., 2017). Hydrophobins are typically found on the outer cell wall surface of aerial hyphae, fruiting bodies and spores, where they reduce wettability, or significantly decrease the surface tension of moist environments, allowing these structures to grow in the air (Wösten et al., 1999). Other roles related to surface perception, attachment to hydrophobic surfaces, and plant colonization have also been shown (Kim et al., 2005; Talbot et al., 1993, 1996). So far, the function of only one *C. fulvum* hydrophobin, HCf-1 (Class I), has been determined. HCf-1 is required for efficient water-mediated dispersal of conidia (Whiteford and Spanu, 2001).

Unlike those described above, BLAST homology searches revealed that most *C. fulvum* ipiSSPs (61 of 70) are novel or have homology to proteins of unknown function. Remarkably, 10 of these ipiSSPs were consistently predicted to have structural homology to proteins present in the RCSB PDB. Of these, CfCE5, CfCE25 and CfCE65 were predicted to be structurally homologous to Alt a 1 from *A. alternata*, which adopts a β−barrel fold unique to fungi (Chruszcz et al., 2012; de Vouge et al., 1996). Recent studies have shown that Alt a 1 is an effector protein with multiple roles in promoting host colonization. Initially, Alt a 1 localizes to the cytoplasm and cell wall of *A. alternata* spores (Garrido-Arandia et al., 2016b; Gómez-Casado et al., 2014). In humid settings, these spores then germinate, and in environments with a pH range of between 5.0 and 6.5, Alt a 1 is released as a tetramer carrying a fungal methoxyflavonol ligand similar to the plant flavonol quercetin (Garrido-Arandia et al., 2016a, b). In the same pH range, which is typical of apoplastic environments, this complex breaks down, releasing Alt a 1 monomers and the flavonol ligand (Garrido-Arandia et al., 2016a,b). The Alt a 1 monomers then function as competitive inhibitors of extracellular plant defence proteins belonging to the pathogenesis-related 5−thaumatin-like protein (PR5-TLP) family (Gómez-Casado et al., 2014), while the flavonol ligand detoxifies reactive oxygen species (ROS) (Garrido-Arandia et al., 2016b). It remains to be determined whether CfCE5, CfCE25 and CfCE65 function in a similar manner during colonization of the tomato leaf apoplast by *C. fulvum*. Interestingly, homologs of CfCE5, CfCE25 and CfCE65 are encoded by the genome of *D. septosporum* (de Wit et al., 2012), and these genes are up-regulated during the infection of pine (Bradshaw et al., 2016). This suggests that the Alt a 1 allergen-like proteins, together with the cerato-platanin, Ecp2-1, Ecp6 and Ecp14-1, which are also ipiSSPs of *D. septosporum* (Bradshaw et al., 2016; de Wit et al., 2012), are core effectors that play important roles in the virulence of both pathogens. These *D. septosporum* ipiSSPs have been shortlisted for future functional characterization (Hunziker et al., 2016).

Four of the nine ipiSSPs, specifically Ecp4, Ecp7, CfCE72 (CTR) and CfCE44, were predicted to be structurally homologous to proteins with a β/γ-crystallin fold. This fold, which typically comprises two four-stranded, anti-parallel Greek key motifs, was originally identified in structural proteins responsible for maintaining the refractive index and transparency of the vertebrate eye lens (Blundell et al., 1981; Wistow et al., 1983). However, this fold is now known to occur in a variety of functionally diverse proteins representing all major taxonomic groups of organisms (Kappé et al., 2010; Mishra et al., 2014). A key feature of this fold in many microbial members is a double clamp N/DN/DXXS/TS Ca^2+^-binding motif required for structure and/or function (Srivastava et al., 2014). This motif, however, is not present in Ecp4, Ecp7, CfCE72 (CTR) or CfCE44.

Strikingly, Ecp4, Ecp7 and CfCE72 (CTR) share a Cys spacing profile with MiAMP1, a plant antimicrobial protein with a β/γ-crystallin fold from nut kernels of *M. integrifolia* (Marcus et al., 1997; McManus et al., 1999). Purified MiAMP1 exhibits broad spectrum inhibitory activity against several plant-pathogenic fungi, oomycetes and gram-positive bacteria *in vitro* (Marcus et al., 1997). Some microbes, however, including several plant- and animal-pathogenic fungi, as well as gram-negative bacteria appear to be insensitive (Marcus et al., 1997). It has been concluded that, to confer broad spectrum antimicrobial activity, MiAMP1 must act on molecules and/or cell structures common to a wide range of microbial organisms (Marcus et al., 1997). Although a specific mode of action for MiAMP1 has not yet been determined (Stephens et al., 2005), more functional information is available for Sp-AMP3, a homolog of this protein from Scots pine, *Pinus sylvestris* (Asiegbu et al., 2003; Sooriyaarachchi et al., 2011). Purified Sp-AMP3 protein has antifungal activity against the plant-pathogenic, root-rotting Basidiomycete *Heterobasidion annosum*, and as part of this, causes morphological changes in the hyphae and spores of this fungus (Sooriyaarachchi et al., 2011). To test the hypothesis that the biological function of Sp-AMP3 involves a fungal cell wall target, carbohydrate-binding assays were performed. These assays revealed that Sp-AMP3 binds to both soluble and insoluble β−1,3-glucans with high affinity, but not to insoluble chitin or chitosan (Sooriyaarachchi et al., 2011). Based on these results, it was hypothesized that differences in cell wall composition would allow Sp-AMP3 to act on some, but not all fungi (Sooriyaarachchi et al., 2011). It is possible that in sensitive fungi, Sp-Amp3 binding interferes with glucan assembly. This could then alter cell wall structure, causing the abovementioned morphological changes, or could result in cell lysis through compromised cell wall integrity (Sooriyaarachchi et al., 2011).

The three remaining ipiSSPs, specifically CfCE24, CfCE56 and CfCE58, were predicted to be structurally homologous to KP6, a killer toxin secreted by specific strains of the fungal corn smut pathogen *U. maydis*. These strains exhibit a “killer” phenotype, which is due to persistent infection by a KP6-producing double-stranded RNA *Totivirus*, P6. Upon secretion, KP6 kills competing, uninfected strains of *U. maydis* (Allen et al., 2013b; Koltin and Day, 1975). Resistance to KP6 in these killer strains is provided by *p6r*, an unknown, non-virus-encoded recessive nuclear host gene (Finkler et al., 1992; Koltin and Day, 1976; Puhalla, 1968). Although a preliminary study suggested that KP6 was only active against grass smut fungi of the order Ustilaginales, with several bacterial and other fungal species shown to be insensitive (Koltin and Day, 1975), it is now clear that KP6 has antifungal activity against other selected plant-pathogenic fungi (Smith and Shah, 2015).

KP6 is translated as a single polypeptide, but is processed into two subunits, KP6α and KP6β, by a kexin protease during passage through the ER−Golgi secretory pathway. This processing involves the removal of a central 31-amino acid residue linker region (Tao et al., 1990), which may serve to keep the two subunits in an inactive protoxin form until the final stages of export (Allen et al., 2013a). Both subunits adopt a core α/β-sandwich fold (Allen et al., 2013a; Li et al., 1999). KP6 functions only as a heterodimer, with both subunits required for cytotoxic activity (Peery et al., 1987). Assays where sensitive *U. maydis* cells were treated with KP6α or KP6β alone, or with one subunit after another, but with a washing step in between, strongly suggest that KP6α is responsible for targeting the cell, while KP6β is cytotoxic (Peery et al., 1987). The specific mode of action for KP6, however, remains unclear. An early study found that spheroplasts derived from a sensitive strain of *U. maydis* were insensitive to KP6, but when the cell wall was given time to regenerate, sensitivity could be restored (Steinlauf et al., 1988). Based on this result, it was inferred that some sort of recognition site was located on the cell wall that then directed KP6 to its cellular target (Steinlauf et al., 1988). However, as was pointed out by Allen et al. (2013b), the cell wall-degrading enzyme preparation used to generate the spheroplasts, Novozyme 234, has residual protease activity (Hamlyn et al., 1981). For this reason, a proteinaceous cell membrane receptor for KP6 cannot yet be ruled out. One possibility is that KP6α forms strong interactions with membrane-associated proteins of the target cell, with KP6β subsequently recruited to the plasma membrane or imported to an intracellular target to cause cell lysis (Allen et al., 2013a). Interestingly, limited amino acid sequence homology was identified between CfCE72 (NTR) and the KP6-like ipiSSPs CfCE24, CfCE56 and CfCE58. This suggests that CfCE72 (NTR) also adopts a KP6-like fold. A putative kexin protease cleavage site is located between the NTR and CTR (β/γ−crystallin-like domain) of CfCE72, implying that this ipiSSP undergoes similar post-translational processing to KP6 upon passage through the ER–Golgi secretory pathway.

In total, 10% of the *C. fulvum* ipiSSPs (seven of 70) are predicted to possess a domain typical of antimicrobial proteins. This raises the possibility that *C. fulvum* dedicates a significant proportion of its apoplastic secretome to functions associated with microbial antagonism, perhaps to outcompete other microbial organisms for nutrients and space in the apoplastic environment, or to provide a form of self-defence (Rovenich et al., 2014). Further studies are now required to establish whether any overlap exists between the *in planta* functions of the β/γ−crystallin/KP6 proteins and the ipiSSPs Ecp4, Ecp7, CfCE24, CfCE44, CfCE56, CfCE58 and CfCE72.

Of course, it remains possible that the predicted similarities in tertiary structure do not extend to biological function. Instead, these folds may be more common than previously thought, irrespective of whether they have evolved from an ancestral protein or by convergent evolution, providing solutions to typical problems faced at the hostile host−pathogen interface. For example, the abovementioned folds may provide enhanced stability in protease-rich environments. Alternatively, they may provide a flexible molecular scaffold for functional diversification and/or the evasion of recognition by corresponding host immune receptors. Recently, the IP effectors Avr1-CO39, AVR-Pia and AvrPiz-t from *M. oryzae*, as well as the ToxB effector from *P. tritici-repentis*, were found to be structurally related (de Guillen et al., 2015). Structure-informed pattern searches subsequently revealed that several other effector candidates from Sordariomycete and Dothideomycete plant pathogens likely share this fold. This led the authors to hypothesize that “the enormous number of sequence−unrelated Ascomycete effectors may in fact belong to a restricted set of structurally conserved effector families” (de Guillen et al., 2015). Certainly, the predicted structural relationship between Alt a 1 and CfCE5/CfCE25/CfCE65 further supports this hypothesis.

Of the 70 apoplastic ipiSSPs from *C. fulvum*, 41 were screened for recognition by wild tomato accessions using an effectoromics approach based on the PVX transient expression system (Hammond-Kosack et al., 1995; Takken et al., 2000). Such an approach has already proven to be successful for the identification of plants carrying immune receptor genes active against other pathogens. For example, of 54 RXLR effectors from the oomycete potato late blight pathogen *Phytophthora infestans*, 31 were found to trigger an HR in one or more of 10 resistant wild *Solanum* accessions, with each accession recognizing between five and 24 effectors (Vleeshouwers et al., 2008). Using the same set of 54 RXLR effectors, 48 were then shown to trigger an HR in one or more of 42 accessions of pepper (*Capsicum annuum*), a non-host of *P. infestans*, with each accession recognizing between one and 36 effectors (Lee et al., 2014). In the current study, nine *C. fulvum* ipiSSPs (Ecps) were found to trigger an HR in one or more of 14 specific wild accessions of tomato. This suggests that nine new IP effectors of this fungus, as well as nine new corresponding *Cf* immune receptor genes, have been uncovered. One of the recognized Ecps, Ecp11-1, is a homolog of AvrLm3, an IP effector from *L. maculans* (Plissonneau et al., 2016). This suggests that both tomato and Brassica carry an immune receptor capable of recognizing this class of effector.

Consistent with *Ecp1, Ecp2-1, Ecp4* and *Ecp5* (Stergiopoulos et al., 2007), but in contrast to *Avr2, Avr4, Avr4E, Avr5* and *Avr9* (Iida et al., 2015; Mesarich et al., 2014; Stergiopoulos et al., 2007), all new *Ecp* genes were found to exhibit limited allelic variation across strains collected from around the world. As has been suggested for *Ecp1, Ecp2-1, Ecp4* and *Ecp5* (Stergiopoulos et al., 2007), this limited allelic variation could reflect a lack of selection pressure imposed on the pathogen to overcome *Cf-Ecp* immune receptor-mediated resistance, since, as far as we are aware, none of the putative corresponding *Cf* immune receptor genes have yet been deployed in commercial tomato cultivars. Alternatively, this lack of allelic variation could reflect selective constraints on the Ecps to maintain their protein sequences (i.e. to ensure full virulence of the pathogen). Of note, all new *Ecp* genes, with the exception of *Ecp14-1*, are associated with repetitive elements in the genome of *C. fulvum* strain 0WU. It is possible that homologous recombination between flanking repeat elements could result in the deletion of these genes, like that hypothesized for strains lacking the repeat-associated *IP* effector genes *Avr4E, Avr5* or *Avr9* (Mesarich et al., 2014; van Kan et al., 1991; Westerink et al., 2004). Thus, to increase potential durability, new *Cf* immune receptor genes should be stacked in resistant tomato cultivars.

In our study, we frequently observed that not all five representatives of a given *S. corneliomuelleri, S. peruvianum*, or *S. pimpinellifolium* accession recognized an Ecp effector. This is not surprising, because both *S. corneliomuelleri* and *S. peruvianum* are typically self-incompatible, while *S. pimpinellifolium* is facultatively self-compatible (Peralta and Spooner, 2006). In other words, genetic variation is expected to exist between representatives of accessions from these species, with this variation extending to the presence or absence of corresponding functional *Cf* immune receptor gene alleles. This may explain why CfCE19 (Ecp17) gave such a strong HR in accession CGN 24034 using the toothpick assay (Fig. S4), but no HR in the agroinfiltration assay (i.e. plants lacking a corresponding functional immune receptor gene allele have been missed by chance) (Fig. S15). This may also be true for Ecp9-1 on CGN 15392 (*S. arcanum* [typically self-incompatible]; Fig. S9), Ecp10-1 on CGN 14356 (Fig. S10), Ecp11-1 on CGN 14357 (Fig. S11), and Ecp13 on CGN 14353 (Fig. S12).

*Cf* immune receptor genes present in self-compatible accessions can be easily introgressed into commercial and breeder’s cultivars of *S. lycopersicum* by backcrossing. In cases of incompatibility, it may be possible to avoid the problems associated with barriers to genetic crossing through a more extensive screen of wild tomato germplasm to identify self-compatible species capable of recognizing the Ecps. This strategy has been successful for the identification of wild potato species that recognize the AVRblb1 IP effector of *P. infestans* (Vleeshouwers et al., 2008). Using an effectoromics approach based on the PVX transient expression system, it was initially determined that the wild potato species *Solanum bulbocastanum*, which is not directly sexually compatible with cultivated potato, *Solanum tuberosum*, carries an immune receptor gene, *RB/Rpi-blb1*, corresponding to *AVRblb1* (Vleeshouwers et al., 2008). As direct introgression of *RB/Rpi-blb1* from *S. bulbocastanum* to *S. tuberosum* is not possible, additional screening was carried out to identify wild potato accessions that are both sexually compatible with cultivated potato and that recognise AVRblb1. HR-associated recognition of AVRblb1 was quickly detected in the sexually compatible species *Solanum stoloniferum*, which was subsequently found to carry *Rpi-sto1*, a functional homolog of *RB/Rpi-blb1* (Vleeshouwers et al., 2008). Importantly, in our study, several accessions were found to recognize the same Ecp effectors, suggesting that this approach could be possible in tomato. Further support is provided by the fact that the *Cf-9* and *Cf-4* immune receptor genes are conserved across the *Solanum* genus (Kruijt et al., 2005; Laugé et al., 2000; van der Hoorn et al., 2001b).

The finding that most new HR-eliciting Ecps have homologs in other plant-pathogenic fungal species raises the possibility of cross-species resistance. In support of this possibility, the Cf-4 immune receptor has been shown to recognize homologs of Avr4 from *D. septosporum, P. fijiensis* and *Pseudocercospora fuligena* (black leaf mould disease of tomato) (de Wit et al., 2012; Kohler et al., 2016; Stergiopoulos et al., 2010), while the Cf-Ecp2-1 immune receptor has been shown to recognize homologs of Ecp2-1 from *D. septosporum* and *P. fijiensis* (de Wit et al., 2012; Stergiopoulos et al., 2012). It must be pointed out, however, that the Cf-4 immune receptor does not recognize homologs of Avr4 from *Cercospora apii, Cercospora beticola* and *Cercospora nicotianae* (leaf spot disease of celery, beet and tobacco, respectively) (Mesarich et al., 2016; Stergiopoulos et al., 2012). With this in mind, it is clear that to provide effective resistance in a recipient plant species, the product of any transferred *Cf* immune receptor gene must recognize an epitope (direct recognition) or virulence function (indirect recognition) conserved to both the corresponding *C. fulvum* effector and its homolog from the target fungal pathogen.

## CONCLUSIONS

In this study, proteomics and transcriptome sequencing were used to identify a set of 70 apoplastic ipiSSPs from *C. fulvum*, which is made up of all 11 IP effectors of this fungus, as well as 59 CfCEs. These ipiSSPs provide new insights into how *C. fulvum* promotes colonization of the tomato leaf apoplast. Using an effectoromics approach, nine CfCEs (Ecps) were found to be recognized by specific wild accessions of tomato. These accessions likely carry new *Cf* immune receptor genes available for incorporation into cultivated tomato.

## MATERIALS AND METHODS

### General

In this study, all kits and reagents were used, unless otherwise specified, in accordance with the manufacturer’s instructions.

### *C. fulvum* strains and tomato accessions

*C. fulvum* strains and tomato accessions used in this study are shown in Tables S5 and S3, respectively.

### Isolation of IWF from the leaf apoplast of *C. fulvum*-infected tomato

Four-to five-week-old H-Cf-0 tomato plants were inoculated with strain 0WU, 4, IPO 1979, or IPO 2559 of *C. fulvum* (compatible interactions). For this purpose, conidia preparation, inoculation, and growth conditions were identical to that described by Mesarich et al. (2014). At 10−17 dpi, IWF was harvested from tomato leaves visibly infected with *C. fulvum* using a previously described protocol (de Wit and Spikman, 1982; Joosten, 2012). Leaf debris and fungal material were then removed by centrifugation at 12,000 × *g* and 4°C for 20 min, and the IWF samples stored at ‒20°C until required.

### Preparation of IWF samples for LC−MS/MS analysis

Frozen IWF samples were thawed on ice and any precipitant formed during the freeze-thaw process removed by centrifugation at 12,000 × *g* and 4°C for 20 min. IWF samples were concentrated 3−300× by: (i) pressure filtration at 4°C using an Amicon 8400 series Stirred Cell Ultrafiltration Unit (EMD Millipore) fitted with an Ultracel regenerated cellulose PLAC 1 kDa nominal molecular weight limit (NMWL) ultrafiltration membrane disc (EMD Millipore); (ii) centrifugation at 4,000 × *g* and 4°C in a 3 kDa NMWL Amicon Ultra-15 Centrifugal Filter Unit (EMD Millipore) or a Vivaspin 20 3 kDa molecular weight cut-off (MWCO) Polyethersulfone (PES) ultrafiltration device (GE Healthcare); or (iii) sequential acetone precipitation, as described by May et al. (1996), with final resuspension in 1 ml dH_2_O. Following concentration, IWF samples were transferred to 2 ml LoBind microcentrifuge tubes (Eppendorf), and stored at ‒20°C until required for further processing. When required, frozen IWF samples were thawed on ice and any precipitant formed during the freeze-thaw process removed by centrifugation at 12,000 × *g* and 4°C for 20 min. A filter-aided sample preparation protocol (Lu et al., 2011), or an in-gel digestion protocol (Karimi Jashni et al., 2015), both based on trypsin digestion, were then used to prepare samples for LC−MS/MS analysis.

### LC–MS/MS analysis

IWF samples were analysed by nano-scale (n)LC−MS/MS with a Proxeon EASY nLC system connected to a LTQ-Orbitrap XL mass spectrometer (Lu et al., 2011) at the Laboratory of Biochemistry, Wageningen University. LC–MS runs and associated MS/MS spectra were analysed with the MaxQuant v1.3.0.5 suite (Cox and Mann, 2008), with default settings applied to the integrated Andromeda peptide search engine (Cox et al., 2011), bar one exception: extra variable modifications were set for the de-amidation of Asn and Gln. MS/MS spectra were searched against one of four sequence databases. These were built from: (i) a collection of common contaminants including, for example, BSA (P02769; bovine serum albumin precursor), trypsin (P00760; bovine), trypsin (P00761; porcine), keratin K22E (P35908; human), keratin K1C9 (P35527; human), keratin K2C1 (P04264; human) and keratin K1CI (P35527; human); (ii) a six-frame translation of tomato (*S. lycopersicum* cv. H-Cf-0) genome sequence (Tomato Genome Consortium, 2012); (iii) the predicted protein catalogue of *C. fulvum* strain 0WU (de Wit et al., 2012; Mesarich et al., 2014), as well as a six-frame translation of the most highly abundant *de novo*-assembled *in vitro* and *in planta* RNA-Seq reads of this fungus (Mesarich et al., 2014); and (iv) a six-frame translation of the repeat-masked *C. fulvum* strain 0WU genome sequence (de Wit et al., 2012). The “label-free quantification (LFQ)” and “match between runs” (set to 2 min) options were enabled. De-amidated peptides were allowed to be used for protein quantification. All other quantification settings were kept at default. Filtering and further bioinformatic analysis of the MaxQuant/Andromeda workflow output and the analysis of abundances for the identified proteins were performed with the Perseus v1.3.0.4 module as part of the MaxQuant suite. Peptides and proteins with a false discovery rate of less than 1%, as well as proteins with at least one peptide across two or more IWF samples, or two or more independent peptides in a single IWF sample, were considered as reliable identification. Reversed hits were deleted from the MaxQuant results table, as were tomato and contamination hits.

### Identification of apoplastic ipiSSPs from *C. fulvum*

*C. fulvum* SSPs directed to the apoplastic environment via the classical/conventional secretory pathway (i.e. SSPs that possess an N-terminal signal peptide, but lack a GPI anchor modification site, a transmembrane domain, or a putative C-terminal ER retention/retention-like signal) were targeted for identification in the protein set identified by LC−MS/MS analysis. The SignalP v3.0 (Bendtsen et al., 2004) and v4.1 (Petersen et al., 2011) servers were used for signal peptide prediction, while the big-PI Fungal Predictor (Eisenhaber et al., 2004) and TMHMM v2.0 (Krogh et al., 2001) servers were used for the prediction of GPI anchor modification sites and transmembrane domains, respectively.

Pre-existing RNA-Seq transcriptome sequencing data (Mesarich et al., 2014) from a compatible *in planta* time course involving *C. fulvum* strain 0WU and *S. lycopersicum* cv. H-Cf-0 (4, 8 and 12 dpi), as well as from strain 0WU grown *in vitro* in PDB or Gamborg B5 liquid media (4 dpi), were used to predict which of the SSPs identified by LC−MS/MS analysis are encoded by *in planta*-induced genes. Although these data lack biological replicates, they have been extensively validated through RT-qrtPCR experiments (Mesarich et al., 2014; this study). Paired-end RNA-Seq reads were re-mapped to the strain 0WU genome sequence (de Wit et al., 2012) with Bowtie v2-2.1.0 (Langmead and Salzberg, 2012) and TopHat v2.0.12 (Kim et al., 2013) using a custom script (Methods S1). Transcript assembly and abundance estimations were then performed using Cufflinks v2.0.2 (Trapnell et al., 2010), with transcript abundance expressed as FPKM values. SSPs were deemed to be *in planta*-induced if they were encoded by genes that had a maximum *in planta* FPKM value of ≥50 at 4, 8 or 12 dpi that exceeded their maximum *in vitro* FPKM value at 4 dpi by a factor of ≥1.5. Gene exon–intron boundaries were confirmed using the same RNA-Seq data.

### General homology screening and alignments

Reciprocal BLASTp screens (Altschul et al., 1997) were used to identify homologs of the apoplastic ipiSSPs from *C. fulvum* present in publicly available databases at NCBI and JGI (Grigoriev et al., 2011). In all cases, hits with an expect (E)-value of >1E-02 were not considered. Likewise, proteins that did not have the same number of Cys residues as the query sequence were not considered. For those proteins for which a homolog could not be identified in JGI, a tBLASTn screen was carried out against the genome collection with the same E-value cut-off. Homologous proteins were aligned using the Clustal Omega server (Sievers et al., 2011).

### Motif identification

The MEME v4.11.2 server (Bailey et al., 2006) was used to identify short sequence motifs shared between members of the *C. fulvum* apoplastic ipiSSP set. For this purpose, the expected distribution of motif sites was set to any number of repetitions per sequence, the number of motifs to find was set to 100, the minimum and maximum length of motif was set to four and 10 amino acid residues, respectively, the minimum and maximum number of sites per motif was set to five and 100, respectively, and the location of motif sites was set to given strand only. All other settings were kept as default.

### Structural modelling

Three-dimensional protein structure prediction servers were used to infer possible structural relationships between apoplastic ipiSSPs of *C. fulvum* and proteins with characterized tertiary structures in the RCSB PDB (Berman et al., 2000). Only those ipiSSPs with no homology to proteins present in NCBI or JGI, or those with homology to hypothetical proteins of unknown function in these databases, were investigated. The prediction servers employed were HHPred (Hildebrand et al., 2009; Söding et al., 2005), SPARKS-X (Yang et al., 2011), MUSTER (Wu and Zhang, 2008), FFAS03/FFAS-3D (Jaroszewski et al., 2005; Xu et al., 2013), FUGUE v2.0 (Shi et al., 2001), RaptorX (Källberg et al., 2012), pGenTHREADER (Lobley et al., 2009), Phyre2 (Kelley et al., 2015) and I-TASSER (Zhang, 2008). Structural modelling was done with MODELLER (HHPred) (Webb and Sali, 2002) and RaptorX, and was visualized using PyMOL (DeLano, 2002). For each server, default settings were used.

### Repeat identification

BLASTn was used to identify repetitive nucleotide sequences shared between the genome scaffolds harbouring an *Ecp* gene and the rest of the *C. fulvum* strain 0WU genome. Only those sequence repeats of ≥100 nucleotides in length, and sharing ≥80% identity, with an E-value threshold of 1E-05, were considered. The maximum total number of sequence alignments considered per scaffold was set to 5,000.

### Homology screening and expression profiling in *Dothistroma septosporum*

Reciprocal BLASTp and tBLASTn screens were used to identify homologs of the apoplastic ipiSSPs from *C. fulvum* in the *D. septosporum* strain NZE10 protein catalogue and genome (de Wit et al., 2012) at JGI, respectively, with hits possessing an E-value of >1E-02 not considered. RNA-Seq data from *D. septosporum* strain NZE10 (Bradshaw et al., 2016) were used to determine which of the homologs are most relevant to the *D. septosporum*−*Pinus radiata* interaction. More specifically, transcript abundance data from one *in vitro* growth condition (fungal mycelia [FM] in *Dothistroma* liquid medium) and three *in planta* growth conditions (epiphytic/biotrophic [early], initial necrosis [mid] and mature sporulating lesion [late]), expressed as reads per million per kb (RPMK) values, were used. Genes deemed relevant to the interaction had to have a maximum *in planta* RNA-Seq RPMK value of ≥50 at the early, mid, or late time point. Furthermore, this value had to exceed the gene’s *in vitro* RPMK value by a factor of at least 1.5.

### PVX-mediated transient expression assays

Tomato accessions (Table S3) were screened for their ability to recognize apoplastic ipiSSPs through the elicitation of an HR using the PVX-based transient expression system (Hammond-Kosack et al., 1995; Takken et al., 2000). For this purpose, the cDNA sequence encoding a mature ipiSSP was fused downstream of the cDNA sequence encoding the *N. tabacum* PR1A signal peptide (i.e. for secretion into the apoplastic environment), and cloned into the binary PVX vector pSfinx behind the *Cauliflower mosaic virus* (CaMV) 35S promoter (Takken et al., 2000). These steps were carried out using the protocol of Mesarich et al. (2014) (overlap extension PCR and restriction enzyme-mediated cloning) or Mesarich et al. (2016) (overlap extension PCR and GATEWAY cloning [Invitrogen]) with the primer pairs listed in Table S6. Constructs were transformed into *Agrobacterium tumefaciens* strain GV3101 for agroinfection of tomato by electroporation using the method of Takken et al. (2000). For localized transient expression assays in tomato, transformants were prepared using the protocol described by Stergiopoulos et al. (2010), but with re-suspension in a final volume of 0.5 ml MMA-acetosyringone, and inoculated into fully expanded leaves by localized wounding on each side of the main vein with a toothpick (Luderer et al., 2002a; Takken et al., 2000). For systemic transient expression assays, transformants were again prepared using the method of Stergiopoulos et al. (2010), with final resuspension in MMA-acetosyringone to an OD_600_ of 1.0, and infiltrated into both cotyledons of a seedling at 10 d post-germination with a 1-ml needleless syringe (Mesarich et al., 2014). The presence or absence of an HR was visually assessed at 10 d post-wounding and 3 weeks post-infiltration for systemic and localized transient expression assays, respectively.

### Tomato infection assays

Tomato accessions (Table S3) were inoculated with *C. fulvum* strain 2.4.5.9.11 IPO using the method described by Mesarich et al. (2014), with resistance or susceptibility to this strain visually assessed across three independent plants at 14 dpi.

### RT-qrtPCR gene expression analysis

Leaf samples from compatible *C. fulvum* strain 0WU–*S. lycopersicum* cv. H-Cf-0 interactions at 4, 8, 12 and 16 dpi, as well as fungal samples from *C. fulvum* strain 0WU PDB and Gamborg B5 liquid media cultures at 4 dpi, were collected by Mesarich et al. (2014) and stored at ‒80°C. Total RNA extraction from each sample, as well as subsequent cDNA synthesis, was carried out according to the protocol of Griffiths et al. (2017). RT-qrtPCR experiments were performed on cDNA samples using the method described by Ökmen et al. (2013) and the primers listed in Table S6. The *C. fulvum actin* gene was targeted as a reference for normalization of gene expression, and results were analysed according to the 2− Δ*C*t method (Livak and Schmittgen, 2001). Results were the average of three biological replicates.

### Allelic variation analysis

*C. fulvum* strains (Table S5) were grown in PDB, with conidia preparation, PDB inoculation, and culture conditions identical to that described by Mesarich et al. (2014). Genomic DNA was extracted from each strain according to the method of van Kan et al. (1991). Genes targeted for an analysis of allelic variation were amplified from genomic DNA by PCR using the protocol and reagents described by Mesarich et al. (2014), and the primers listed in Table S6. PCR amplicons were purified using an illustra GFX PCR DNA and Gel Band Purification Kit (GE Healthcare), and were directly sequenced at Macrogen Inc. (Korea) using the same gene-specific primers employed for PCR amplification.

## ACKNOWLEDGEMENTS

We thank Simon Williams (Australian National University, Canberra, Australia) for advice on protein structure prediction, Andre Sim (Massey University, Palmerston North) for mapping RNA-Seq reads to the *D. septosporum* genome, Sjef Boeren (Wageningen University, the Netherlands) for performing LC−MS/MS experiments, and Willem van Dooijeweert (Centre for Genetic Resources, the Netherlands) for providing tomato seed. Financial assistance for this research was provided by Wageningen University, the Royal Netherlands Academy of Arts and Sciences, European Research Area−Plant Genomics, the Centre for BioSystems Genomics (part of the Netherlands Genomics Initiative/Netherlands Organization for Scientific Research; project TD8-35), and the New Zealand Bio-Protection Research Centre. Financial assistance for CW was provided by the Chinese Scholarship Council. The authors declare no conflicts of interest.

## AUTHOR CONTRIBUTIONS

CHM, BÖ, REB, MDT and PJGMdW conceived the project. CHM, CHD and AvdB performed the bioinformatic analyses. CHM, BÖ, HR, SAG, CW, MKJ, AM, JC, LH and HGB carried out the experimental work. CHM wrote the manuscript. All authors read and approved the final manuscript.

## AUTHOR-RECOMMENDED INTERNET RESOURCES

Big-PI Fungal Predictor server: http://mendel.imp.ac.at/gpi/fungi_server.html

Clustal Omega server: https://www.ebi.ac.uk/Tools/msa/clustalo/

FFAS03/FFAS-3D server: http://ffas.sanfordburnham.org/ffas-cgi/cgi/ffas.pl

FUGUE v2.0 server: http://mizuguchilab.org/fugue/prfsearch.html

HHPred server: https://toolkit.tuebingen.mpg.de/hhpred/

I-TASSER server: http://zhanglab.ccmb.med.umich.edu/I-TASSER/

JGI BLAST server: http://genome.jgi.doe.gov/pages/blast-query.jsf?db=fungi

MEME v4.11.2 server: http://meme-suite.org/tools/meme

MUSTER server: http://zhanglab.ccmb.med.umich.edu/MUSTER/

NCBI BLAST server: https://blast.ncbi.nlm.nih.gov/Blast.cgi

pGenTHREADER server: http://bioinf.cs.ucl.ac.uk/psipred/

Phyre2 server: http://www.sbg.bio.ic.ac.uk/phyre2/html/page.cgi?id=index

RaptorX server: http://raptorx.uchicago.edu/StructurePrediction/predict/

RCSB PDB: http://www.rcsb.org/pdb/home/home.do

SignalP v3.0 server: http://www.cbs.dtu.dk/services/SignalP-3.0/

SignalP v4.1 server: http://www.cbs.dtu.dk/services/SignalP/

SPARKS-X server: http://sparks-lab.org/yueyang/server/SPARKS-X/

TMHMM v2.0 server: http://www.cbs.dtu.dk/services/TMHMM/

## GENBANK ACCESSION NUMBERS

*Ecp6*, KX943112; *Ecp7*, KX943113; *Ecp8/CfCE6*, KX943038; *Ecp9-1/CfCE9*, KX943041; *Ecp9-2*, KX943114; *Ecp9-3*, KX943115; *Ecp9-4*, KX943116; *Ecp9-5*, KX943117; *Ecp9-6*, KX943118; *Ecp9-7*, KX943119; *Ecp9-8*, KX943120; *Ecp9-9/CfCE49*, KX943081; *Ecp10-1/CfCE14*, KX943046; *Ecp10-2/CfCE31*, KX943063; *Ecp10-3*, KX943121; *Ecp11-1/CfCE18*, KX943050; *Ecp12/CfCE26*, KX943058; *Ecp13/CfCE33*, KX943065; *Ecp14-1/CfCE55*, KX943087; *Ecp14-2*, KX943122; *Ecp15/CfCE59*, KX943091; *Ecp16/CfCE48*, KX943080; *Ecp17/CfCE19*, KX943051; *CfPhiA-1/CfCE11*, KX943043; *CfPhiA-2/CfCE53*, KX943085; *CfCE3*, KX943035; *CfCE4*, KX943036; *CfCE5*, KX943037; *CfCE7*, KX943039; *CfCE8*, KX943040; *CfCE12*, KX943044; *CfCE13*, KX943045; *CfCE15*, KX943047; *CfCE16*, KX943048; *CfCE20*, KX943052; *CfCE22*, KX943054; *CfCE24*, KX943056; *CfCE25*, KX943057; *CfCE27*, KX943059; *CfCE30*, KX943062; *CfCE34*, KX943066; *CfCE35*, KX943067; *CfCE36*, KX943068; *CfCE37*, KX943069; *CfCE40*, KX943072; *CfCE41*, KX943073; *CfCE42*, KX943074; *CfCE44*, KX943076; *CfCE47*, KX943079; *CfCE50*, KX943082; *CfCE51*, KX943083; *CfCE56*, KX943088; *CfCE57*, KX943089; *CfCE58*, KX943090; *CfCE60*, KX943092; *CfCE61*, KX943093; *CfCE63*, KX943094; *CfCE64*, KX943095; *CfCE65*, KX943096; *CfCE66*, KX943097; *CfCE67*, KX943098; *CfCE68*, KX943099; *CfCE69*, KX943100; *CfCE70*, KX943101; *CfCE71*, KX943102; *CfCE72*, KX943103; *CfCE73*, KX943104; *CfCE74*, KX943105; *CfCE76*, KX943107; *CfCE77*, KX943108.

